# Joint mechanistic modeling of viral and antibody responses to vaccines in non-human primates to quantify SARS-CoV-2 mechanistic correlates of protection

**DOI:** 10.64898/2026.01.15.699647

**Authors:** Marie Alexandre, Romain Marlin, Laetitia Bossevot, Mariangela Cavarelli, Nathalie Dereuddre-Bosquet, Francis Relouzat, Mireille Centlivre, Roger Le Grand, Rodolphe Thiébaut, Yves Lévy, Mélanie Prague

## Abstract

In a global SARS-CoV-2 landscape of hybrid immunity resulting from a vaccinated worldwide population and the emergence of new variants able to escape immunity, the identification and quantification of correlates of protection (CoPs) is crucial for the adaptation of next-generation SARS-CoV-2 vaccines. Antibodies and their neutralizing capacity have been identified as reliable mechanistic CoPs. Here, we proposed an original mechanistic model jointly describing viral and antibody dynamics, and their mutual interactions, observed after SARS-CoV-2 infection in non-human primates (NHPs) with distinct immunological backgrounds. From the model, the concentration of neutralizing antibodies being protective against viral infection spreading was derived using the reproduction number. Counterfactual simulations were also performed to better understand and validate immune mechanisms driving immune control. The model was estimated on viral, binding (bAb), and neutralizing (nAb) antibody dynamics, with data collected in 34 naive and convalescent NHPs. Animals were involved in a preclinical study evaluating two next-generation protein-based vaccines targeting the RBD of the Spike protein to CD40-expressing cells, and the original BNT162b2 mRNA vaccine against Delta SARS-CoV-2 infection. Our model validated the functionality of nAb to neutralize viruses as the primary mechanism of protection. An inhibitory antibody concentration against Delta variant of 20 AU/mL was deemed protective against Delta infection in naive animals. Moreover, we showed the strong benefit of hybrid immunity to induce faster and more protective antibody responses pointing out the crucial role of the memory B-cell immune response in viral control. Finally, an additional effect of natural immunity beyond antibodies and enhancing the elimination of infected cells was identified, suggesting the potential role of the T-cell response in viral control. Our model showed the benefit of combining information from both viral and antibody responses to better qualitatively and quantitatively inform on their protective capacity against Delta SARS-CoV-2 infection.

## Introduction

The severe acute respiratory syndrome coronavirus 2 (SARS-CoV-2), which emerged in Wuhan at the end of 2019, is the causative agent of the coronavirus disease 2019 (COVID-19) pandemic. The respiratory disease was responsible for 7.1 million reported deaths worldwide and over 27.3 million including excess mortality in June 2024 [1]. Several vaccines were rapidly developed and approved to combat the spread of the disease and were available for large-scale vaccination campaigns at the end of 2020 [2]. Initially developed using sequence information from the original Wuhan strain of SARS-CoV-2, the first-generation COVID-19 vaccines demonstrated high efficacy in preventing symptomatic and severe COVID-19 [3,4], and in reducing virus transmission [5–7]. However, the emergence of new variants of concern (VoCs) from the end of 2020 and their ability to escape immunity [2,8], significantly modified the worldwide immunological landscape [9]. In particular, the Delta and more recently the Omicron variant and sub-variants led to reduced efficacy of first-generation vaccines and waning immunity [10–12], requiring a complete 2-dose vaccination strategy [13,14] or a booster dose [10,15,16], and inducing an increased transmission even in vaccinated individuals [2,17–19].

Most first-generation COVID-19 vaccines focused on the spike (S) glycoprotein of SARS-CoV-2. The rationale was that the receptor-binding domain (RBD) within S binds to the angiotensin-converting enzyme 2 (ACE2) receptor, located on the surface of host cells, to enter into cells [20]. Antibodies directed against the S protein and specifically against RBD could then prevent this interaction, neutralizing the virus and protecting individual from infection [2,21]. Correlates of protection (CoPs), which are immunological markers that reliably predict protection against infection or symptomatic disease, are a key point in vaccine development [22]. Antibody titers, and more specifically neutralizing antibodies, have been widely studied and identified as CoPs for COVID-19 vaccines against pre-Omicron virus [23–32]. However, the emergence of the Omicron variant, and its rapid antigenic shift, which led to extensive mutations in the spike and RBD, enabled sustained and efficient escape from the neutralizing antibodies induced by the first-generation vaccines [2,33]. Accordingly, it pointed out the necessity to develop next-generation SARS-CoV-2 vaccines providing better protection against emerging VoCs [34] and also tackling effects of immune imprinting [35], which refers to the immune system’s tendency to preferentially recall and respond to antigens it first encountered. Furthermore, in this context of constant and rapid viral evolution, a larger proportion of the population becomes susceptible to infection despite vaccination leading to a landscape of hybrid immunity. Considering that CoPs can differ from immunity induced by vaccination or infection [36], it appears essential to identify and quantify CoPs in a global context of viral evolution. Although identifying a CoP is challenging, the characterization of mechanistic CoP (mCoP), i.e. immune marker that is causally associated with protection, is even more difficult. In particular, it requires the ability to experimentally control immunization events and longitudinally collect multiple immune markers to directly be able to evaluate mechanistic causation of immune responses [23].

As pointed out by Schiffer [37], preclinical studies, that provide the opportunity to evaluate, early after the challenge, viral and immune responses in controlled infection models, and mathematical modeling are undoubtedly fundamental tools in this process. To study respiratory tract infections, non-human primate (NHP) models are particularly relevant for the development of vaccines. Indeed, NHPs share anatomically and physiological similarity to humans, and display similar innate and adaptive immune responses against viral infections. Moreover, closed similarities between human and NHP ACE2 receptors accentuate the benefit of animal models in the case of SARS-CoV-2 [38].

In a previous work [32], we proposed a mechanistic model-based approach applied on NHP preclinical studies to evaluate the immune mechanisms involved in vaccine- and natural-induced immunity, and identify mechanisms of protection and immune markers capturing them, i.e., mCoPs. More specifically, this work allowed us to identify neutralizing antibodies inhibiting the binding between ACE2 receptor and RBD as reliable mCoP able to capture effects elicited by immunization to reduce the ability of SARS-CoV-2 virions to infect target cells. An additional effect of natural immunity on an enhanced elimination of infected cells was found but without identification of mCoP. The next step consists now to quantify the mCoP by defining a protective threshold against infection. To this end, we proposed to build a mechanistic model based on ordinary differential equations (ODEs) jointly describing the within-host interactions of the virus with its host, the humoral responses induced after infection in immunized or naive individuals, and their mutual interactions. As highlighted by two recent systematic reviews [33,39], only a limited number of mechanistic models have been proposed in the literature to simultaneously describe viral and humoral responses in infectious diseases. Numerous models have been developed to describe viral dynamics following infection alone, or including treatment effects, mostly based on target-cell-like models [40–45]. A few models, more or less complex, have been proposed to describe humoral responses and antibody dynamics following vaccination [46–49]. Among models including both viral and immune responses, some only focused on the impact of innate immune responses on viral dynamics [50–53], mostly through interferon dynamics, while other integrated the full immune response (i.e., T-cells, B-cells, antibody, memory response, cytokines), but were too complex to be estimated on data and consequently were only developed for a theoretical purpose [54–57]. Nevertheless, three models were still found in the literature [58–60] combining both viral and antibody dynamics and estimated on data, and helped us to build our model. We applied our modeling work to a preclinical study recently published [61] assessing efficacy of two next-generation protein-based vaccines targeting the RBD of the SARS-CoV-2 spike protein to CD40-expressing antigen-presenting cells. We have developed an antibody-mediated targeting vaccine (AMV) platform, utilizing a fully humanized monoclonal antibody targeting selected pathogen epitopes to the CD40 receptor, thereby enhancing antigen delivery and dendritic cell stimulation. We designed a bivalent CD40 construct presenting both the RBD from the original Wuhan strain and a variant RBD incorporating K417N, E484K, and N501Y mutations (RBDv1). To broaden the scope of SARS-CoV-2 vaccines beyond the S protein, a second candidate, CD40.Pan.CoV, includes a highly conserved non-S Nucleocapsid sequence (Npep2) and an RBD sequence harboring VoC mutations[61]. Vaccines were proven efficient to boost protective immune responses against SARS-CoV-2 in convalescent NHPs and are currently tested in phase 1/2a clinical trials. In this work, we first validated neutralizing antibodies as robust mCoP before identifying immune mechanisms of actions explaining the distinct antibody dynamics observed according to NHP immunological backgrounds. Finally, a protective level of neutralizing antibody allowing rapid control of viral replication following SARS-CoV-2 new infection was derived.

## Materials and methods

### Ethics statement

All animals were housed in IDMIT facilities (CEA, Fontenay-aux-roses, France), under BSL-2 and BSL-3 containment when necessary (Animal facility authorization #D92-032-02, Préfecture des Hauts de Seine, France) and in compliance with European Directive 2010/63/EU, the French regulations and the Standards for Human Care and Use of Laboratory Animals, of the Office for Laboratory Animal Welfare (OLAW, assurance number #A5826-01, US). The protocols were approved by the institutional ethical committee ‘Comité d’Ethique en Expérimentation Animale du Commissariat à l’Energie Atomique et aux Energies Alternatives’ (CEtEA #44) under statement number A20-061.The study was authorized by the ‘Research, Innovation and Education Ministry’ under registration number APAFIS#28946-2021011312169043 v2.

### Data

#### Experimental procedure

Data comes from a study [61] performed on cynomolgus macaques (*Macaca fascicularis*) evaluating the efficacy of two protein-based vaccines targeting the RBD of the SARS-CoV-2 spike protein to CD40 and one mRNA vaccine to boost protective immunity in SARS-CoV-2 convalescent NHPs. The CD40.RBDv vaccine is a humanized anti-human CD40 12E12 IgG4 monoclonal antibody fused via the heavy chain to SARS-CoV-2 RBD from the original Wuhan strain (2019-nCoV), and via the light chain to the SARS-CoV-2 RBDv1 incorporating mutations described in VoCs Alpha, Beta, Gamma and Omicron subvariants (K417N, E484K, and N501Y). The CD40.Pan.CoV vaccine is a humanized anti-human CD40 12E12 IgG4 monoclonal antibody fused via the C-terminal of the heavy chain to SARS-CoV-2 Npep2, and of the light chain to SARS-CoV-2 RBDv2 harboring mutations described in VoCs Alpha, Beta, Gamma, Delta, Kappa and Omicron subvariants (K417N, L452R, T478K, E484Q, N501Y). The BNT162b2 mRNA vaccine (Pfizer-BioNTech) is a lipid nanoparticle–formulated, nucleoside-modified RNA vaccine that encodes a prefusion stabilized, membrane-anchored Wuhan SARS-CoV-2 full-length spike protein.

The study includes 34 macaques, aged from 37 to 76 months, and including 20 females and 14 males. Among them, 23 animals were first exposed to a high dose (i.e. 1 × 10^6^pfu) of historical SARS-CoV-2 strain, administrated via the intra-nasal and intra-tracheal routes simultaneously. As shown in Figure 1, after an average convalescent phase of 18.2 months (mo) (median=17.7 mo; IQR = [16.5, 17.7] mo) (Table S1), NHPs were randomized to receive either a placebo (n=6, referred to as Convalescent group), a subcutaneous injection of CD40.RBDv vaccine (n=6, referred to as CD40.RBDv-Conv group), a subcutaneous injection of CD40.Pan.CoV vaccine (n=5, referred to as CD40.PanCoV-Conv group), or an intramuscular injection of monovalent mRNA Bnt162b2 vaccine (Pfizer) (n=6, referred to as mRNA-Conv group). Two additional groups of naive animals were included receiving either a placebo (n=5, referred to as Naive group), or an injection of CD40.RBDv vaccine (n=6, referred to as CD40.RBDv-Naive group). Four weeks after vaccination, all macaques were challenged with a high dose (1 × 10^5^ TCID_50_) of B.1.617.2 SARS-CoV-2 (also known as the Delta variant). Over the 5 mL of virus preparation, 0.5 mL were inoculated via the intra-nasal route (0.25 mL in each nostril), and 4.5 mL via the intra-tracheal route.

**Figure 1.**
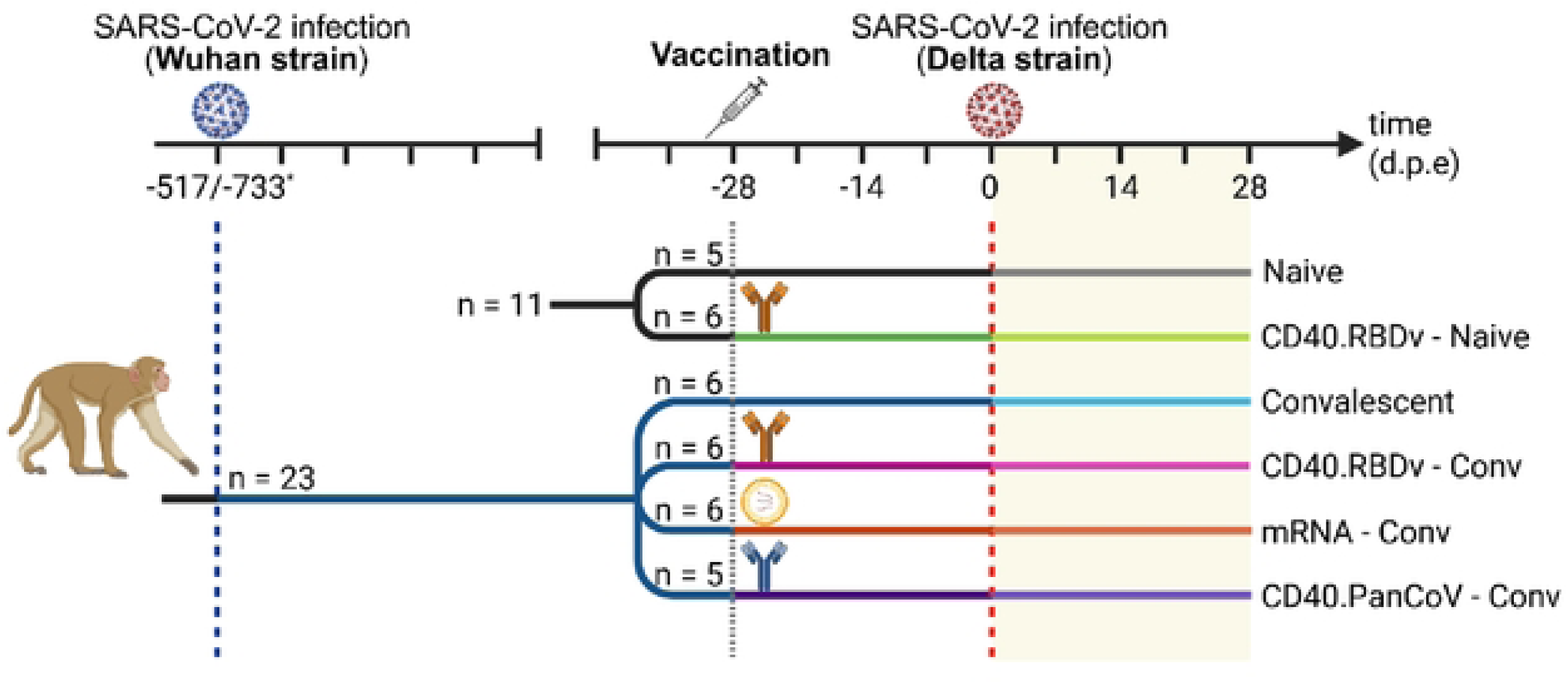
Study design. Schematic overview of immunization (infections and vaccinations) strategies in 34 naive and Wuhan SARS-CoV-2 convalescent animals. In naïve animals (n=11, in black), cynomolgus macaques were injected subcutaneously (SC) with CD40.RBDv (n=6, light green) or included as controls (n=5, gray). In Wuhan convalescent animals (n=23, dark blue), macaques were injected SC with CD40.RBDv (n=6, pink), CD40.PanCoV (n=5, purple), intramuscularly with BNT162b2 mRNA vaccine (Pfizer-BioNTech; n=6, orange) or included as controls (n=6, light blue). Four weeks after vaccination (day 0), all animals were exposed to a high dose of B.1.617.2 SARS-CoV-2 (Delta strain) and were longitudinally followed for 30 days post-exposure (d.p.e), with regular collection of nasopharyngeal and tracheal fluids, and blood samples. Only data collected post Delta SARS-CoV-2 infection (yellow area) were included in the modeling. The vertical lines represent the initial Wuhan SARS-CoV-2 infection (blue dashed line), the vaccination (gray dotted line) and the Delta SARS-CoV-2 infection (red dashed line).

#### Evaluation of viral and antibody responses

For each animal, nasopharyngeal and tracheal fluids were collected at 0, 1, 2, 2, 3, 4, 5, 7, 8, 14, 23 and 29 days post-exposure (d.p.e), and blood samples for measuring humoral immune responses were collected at 0, 3, 9, 13 and 22 d.p.e. Both viral genomic RNA (gRNA) and subgenomic RNA (sgRNA) were quantified in nasopharyngeal and tracheal swabs by the RT-qPCR targeting RdRp-IP4 and SARS-CoV-2 E gene leader sequences, respectively (Figure S1).

Two types of humoral immune responses were measured in blood: 1) anti-RBD binding IgGs titrated using a commercially available multiplexed immunoassay developed by Mesoscale Discovery (MSD, Rockville, MD) and expressed as arbitrary unit (AU)/mL, 2) antibodies neutralizing the binding of the RBD domain to the ACE2 receptor titrated using the MSD pseudo-neutralization assay and expressed in AU/mL (Figure S2). In these two MSD assays, serology kits detecting especially antibodies that target antigens from SARS-CoV-2 Delta variants were used.

### Statistical analysis

A preliminary descriptive analysis was performed on the viral and antibody dynamics observed in the 34 NHPs to describe, summarize and compare dynamics between groups. In particular, the viral dynamics were characterized by the following metrics: 1) the time to and value of peak viral load, 2) the area under the viral load curve (AUC), 3) the duration of the acute stage, defined as the time interval between the first and the last detectable viral load, and 4) the duration of the clearance stage, defined as the time interval between the peak and the first undetectable viral load. In turn, antibody dynamics were characterized by: 1) the area under the immune curve, and 2) the value at exposure. Statistical differences among groups of animals were evaluated using classic two-sample t-tests (Welch’s 𝑡-test in case of unequal variance, identified by a F-test, and Student 𝑡-test otherwise) and global ANOVA tests for peak VL and AUC, while two-sample Mann-Whitney tests and global Kruskal-Wallis tests were used for time-based descriptors. P-values were adjusted for test multiplicity with Benjamini and Hochberg correction [62].

### Joint model for viral and immune responses

We proposed a mechanistic model jointly describing the dynamics of viral and humoral responses following infection in animals, vaccinated or not. In our population approach, the mechanistic model is divided in three layers: (1) a mathematical model based on ordinary differential equations (ODEs) to jointly describe the within-host viral dynamics in the two compartments of the upper respiratory tract (URT), the trachea and nasopharynx, and immune responses, (2) a statistical model to account for inter-individual variability and the effects of immunological background on individual dynamics, and (3) an observation model linking observed viral and antibody dynamics to ODE compartments.

### Mechanistic model

#### Mathematical model for viral dynamics

We used a mechanistic model previously published [32] to describe SARS-CoV-2 viral gRNA and sgRNA dynamics in a similar NHP vaccine study [63]. In each URT compartment, a target-cell limited model-like is considered assuming that after infection, uninfected target cells (𝑇) can be infected (𝐼_1_) by inoculated viruses (𝑉_𝑠_) at an infectivity rate 𝛽 = 𝛽_0_(1 ― 𝜀_𝐴𝑏_). This rate is defined as the product of two terms: 𝛽_0_ being the constant viral infectivity rate in the absence of an effective immune response, and a second term, (1 ― 𝜀_𝐴𝑏_), characterizing the effect of the immune response to block infection spreading and that is fully defined below. After an eclipse phase of 1/𝑘 days, infected cells become productively infected cells (𝐼_2_) and can produce virions at a rate 𝑃, of which proportions 𝜇 and (1 ― 𝜇) are infectious (𝑉_𝑖_) and non-infectious (𝑉_𝑛𝑖_), respectively. After their production, *de novo* produced virions can in turn infect target cells. Productively infected cells, *de novo* produced virions and inoculated virions are cleared at rates 𝛿, 𝑐 and 𝑐_𝑖_, respectively. Accordingly, the viral model is described through Equations (1) to (6), where the superscript 𝑋 denotes the URT compartment (N: nasopharynx; T: trachea). As described in the literature [41,45] and demonstrated in our previous work [32], viral transport between URT compartments is negligible compared to viral clearance in viral kinetics. Accordingly, no virus exchange between the trachea and nasopharynx was considered in this model. We defined Ψ_𝑣𝑙_ = (𝛽,𝛿,𝑃^𝑇^,𝑃^𝑁^,𝜇,𝑘,𝑐,𝑐)^𝑇^ the vector of model parameters characterizing the viral dynamics.

#### Mathematical model for antibody dynamics

To take into account the effects of the immune response on viral dynamics after infection, we proposed a model integrating an antigen-mediated immune response. In particular, this modeling of the immune response aims at considering the immunological background of animals to explain their distinct ability of viral control, whether partial or total, after infection despite their similar challenge. Based on our previous work [32], we focused especially on the role of antibodies in blocking new infections. Modeling the dynamics of the humoral immune cells only on a short period of 30 days following infection, we proposed a simple model including two types of cells, antibody secreting cells (ASCs) and antibodies. After infection, *de novo* produced virions, whether infectious or non-infectious, trigger the humoral immune response resulting in the generation of ASCs (𝑆) at rate 𝛾, and that are eliminated at rate 𝛿_𝑆_. This compartment accounts for all populations of plasma cells able to produce antigen-specific antibodies. After the establishment of the immune response and viral clearance, antibody level remains stable in time [60]. This stabilization mostly results in the humoral response driven by long-lived plasma cells predominantly produced by germinal center (GC)-derived plasma cells [49], and by the memory B cells. To include this homeostasis in a simple way, a logistic function was added in the dynamics of 𝑆 [59], with a logistic growth rate 𝜌 characterizing the differentiation rate of B cells (including memory B cells and GC) into ASCs, and a carrying capacity 𝑆_𝑚𝑎𝑥_. Antibodies are produced by 𝑆 at rate 𝜃 and are eliminated at rate 𝛿_𝐴𝑏_. Both binding (bAb) and neutralizing (nAb) antibodies being observed in our data, these two populations have been included in the model, defining nAb as a sub-population of bAb. In other words, we assumed that only a proportion 𝛼_𝑛𝑎𝑏_ ∈ (0,1) of binding antibodies is able to neutralize the virus. The choice of this proportional relationship was directly driven by data (see Figure S3). As previously identified [32], we considered the blockade of new infections as the main immune mechanism driven by the humoral response, and more specifically by neutralizing antibodies. We modeled the ability of nAb to neutralize the virus by reducing the viral infectivity 𝛽_0_ by the factor (1 ― 𝜀_𝐴𝑏_), with the efficacy of neutralization 𝜀_𝐴𝑏_ defined by an Emax function [60]: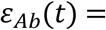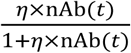, and whose value lies between 0 in the absence of nAb and 1 for high levels of antibodies. The parameter 𝜂 represents the net effect of nAb on controlling viral infection, or 1/𝜂 can be seen as the concentration of nAb, in AU/mL, required to reach 50% of neutralization efficacy. We defined 𝚿_𝐴𝑏_ = (𝛾,𝜌,𝑆_𝑚𝑎𝑥_,𝛿_𝑆_,𝜃,𝛿_𝐴𝑏_,𝛼_𝑛𝑎𝑏_,𝜂,𝑆_0_)^𝑇^ the vector of model parameters describing the humoral response. The full model we built is given in the following set of Equations (1) to (9) and its schematic diagram is presented in Figure 2.

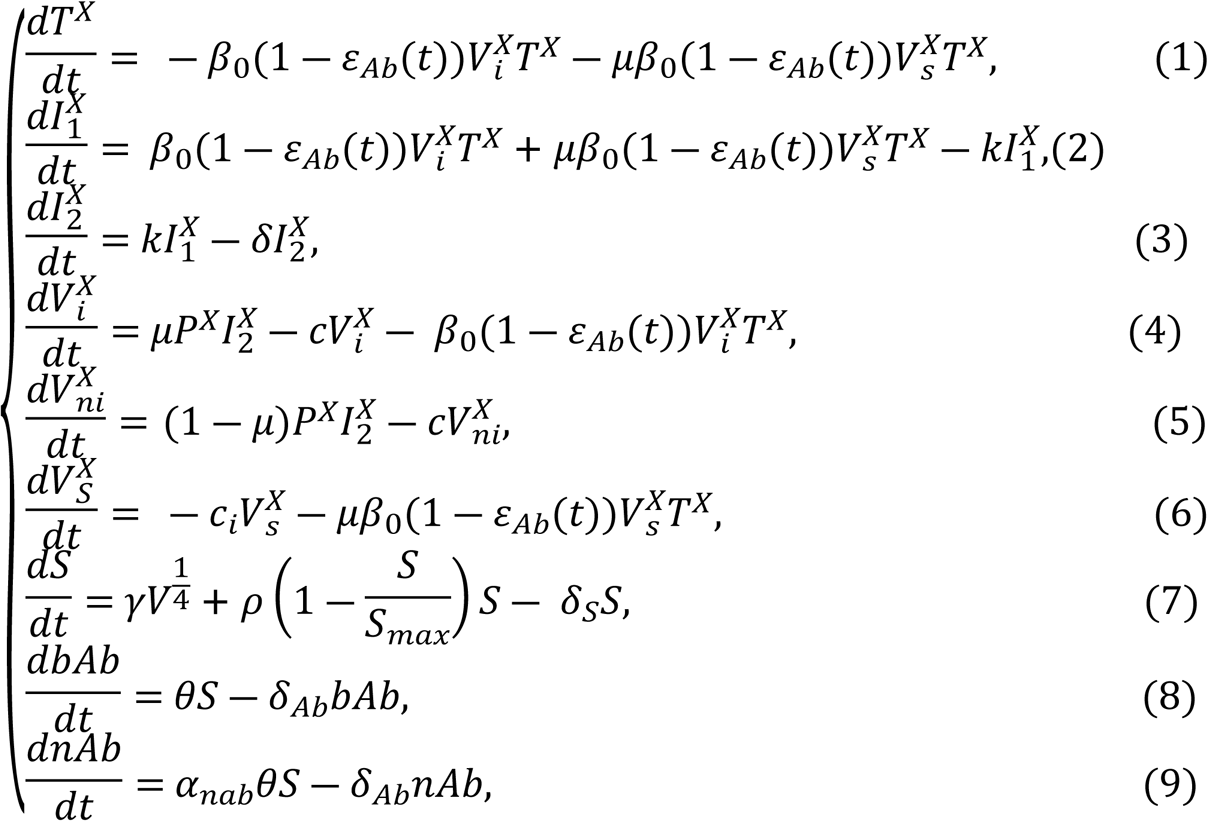

where 𝑉 represents the total concentration of free virions produce in the two URT compartments, such that 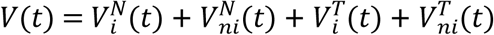. To model the triggering of the humoral response by the presence of antigens, we initially assumed a production of 𝑆 proportional to 𝑉. However, to overcome identifiability issues on the parameter 𝛾 resulting from the difference of scales between the viral and humoral dynamics in our data, several classic transformations of 𝑉 have been tested, as log-, root- or fourth root-transformations. The model using the fourth root transformation has proven to better improve data fitting (i.e. being the model with the best BICc value) explaining the production term of 𝑆 proportional to 𝑉^1/4^ in Equation (7). Although more complex models have already been proposed to describe antibody dynamics after antigenic stimulation [33,46–49], no distinction has been made here between short-lived and long-lived ASCs, two populations of plasma cells differing by their life expectancy, because of short time of follow-up and the resulting identifiability issues. Mathematically, a single ODE compartment (𝐴𝑏) for antibodies could be considered in our final model, with 𝛼_𝑛𝑎𝑏_ × 𝐴𝑏 and 𝐴𝑏 representing the populations of neutralizing and binding antibodies, respectively. Distinct compartments have been considered here for a better visual understanding. The choice of the non-linear Emax function between the concentration on neutralizing antibodies and their efficacy to neutralize viruses relies on the sigmoidal shape of this relationship observed in our previous work [32] and in other preclinical vaccine trials against SARS-CoV-2 conducted on mice (internal unpublished work). Although we focused our interest in the functionality of antibodies to neutralize viruses after infection, other immune mechanisms have already been mentioned in the literature. In particular, two other potential effects of virus-specific antibodies have been studied [32,59,60,64,65]: first, their ability to enhance virus clearance due to opsonization, and second to stimulate destruction of infected cells through antibody-dependent cellular cytotoxicity or complement-mediated lysis. We defined 𝚿 = 𝚿_𝑣𝑙_ ∪ 𝚿_𝐴𝑏_ the vector of parameters of the full model to be fixed or estimated.

**Figure 2.**
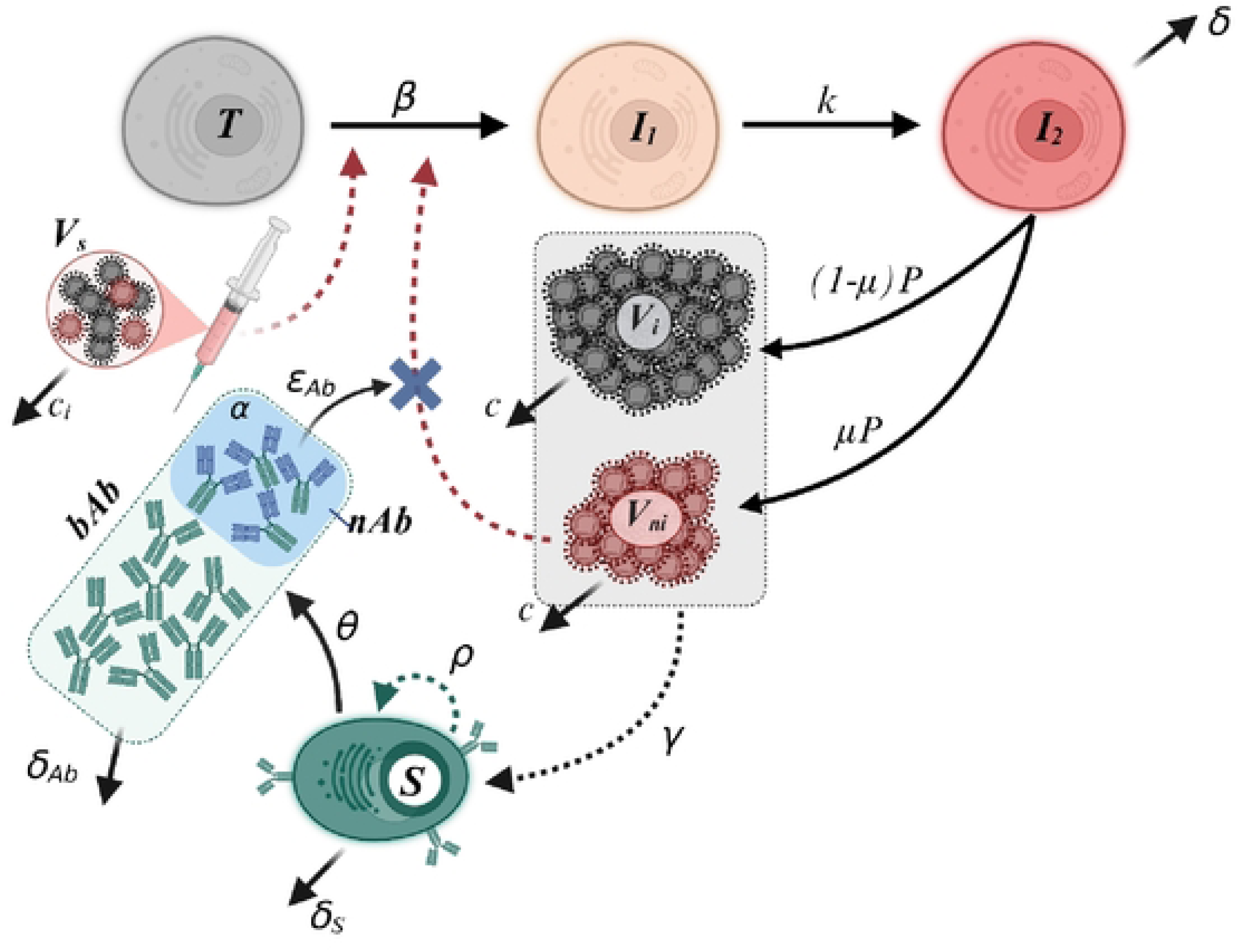
Schematic diagram of the model proposed to jointly describe viral and antibody dynamics following SARS-CoV-2 infection with Delta strain. To simplify visual appearance, no distinction between the two URT compartments (trachea and nasopharynx) has been made for viral dynamics (i.e. for 𝑇, 𝐼_1_, 𝐼_2_, 𝑉_𝑖_, 𝑉_𝑛𝑖_ and 𝑉_𝑠_ entities).

#### Assumption on parameter values

Assuming the time of exposure as the initial time 𝑡 = 0 and that animals were either naive of infection or convalescent of a previous infection occurring several months before, we considered the absence of virions and a full pool of target cells at the time of injection, hence 𝑇 = 𝑇_0_. Moreover, the last stimulation of the immune system occurring at the earliest four weeks before exposure for all NHPs, we assumed that antibody dynamics reached a steady state at the time of infection. Accordingly, the initial conditions of our model are defined as

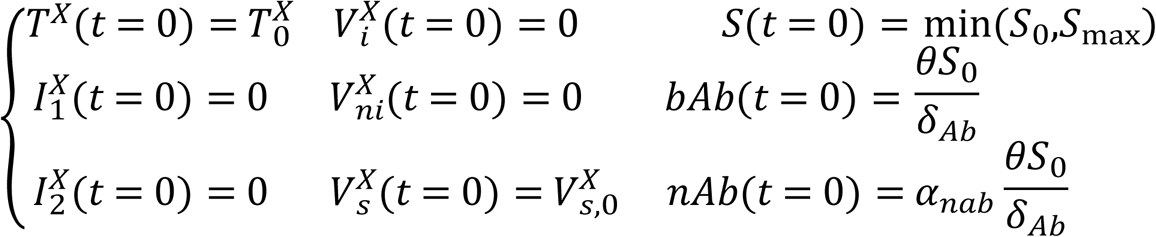

where the initial concentrations of target cells, i.e., epithelial cells expressing the ACE2 receptor, are defined as 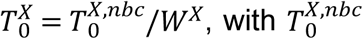 and 𝑊^𝑋^ the initial number of target cells and the volume of distribution of compartment 𝑋. The concentrations of virions inoculated are given by 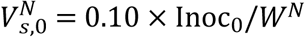 and 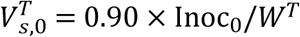, with Inoc_0_ being the total number of virions injected. To overcome the absence of long-lived plasma cells and memory B cells, the immunological background was integrated into the model by adjusting the initial condition of 𝑆 for the groups of treatment (see *Statistical model*). Moreover, this latter was defined as the minimal value between 𝑆_0_and 𝑆_𝑚𝑎𝑥_to meet the condition of saturation of the compartment 𝑆 induced by the logistic function.

Some parameters have been fixed based on experimental settings and/or literature to ensure the identifiability of the model (see Table 1). First, the initial numbers of target cells were fixed to 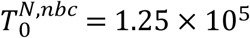 cells and 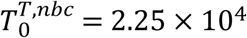 cells in the nasopharynx and the trachea, respectively [41]. Similarly, the number of inoculated virions was experimentally fixed to Inoc_0_ = 1.40 × 10^9^ virions, resulting from an inoculation of 10^5^TCID50 with a concentration of virions of 2.80 × 10^8^ virions/mL in a sample of 5mL. Based on our previous modeling work [32], the eclipse rate was fixed to 𝑘 = 3 day^―1^, corresponding to a mean duration of the eclipse phase of 8 hours, and the ratio between infectious and non-infectious virions was set to 𝜇 = 10^―3^ (Marc et al. [65] demonstrated the absence of effect of Delta variant on this parameter compared to the historical strain). In addition, the viral clearances of inoculated and *de novo* produced virions were fixed to 𝑐_𝑖_ = 20 day^―1^ and 𝑐 = 3 day^―1^, respectively. A profile likelihood was performed to re-calibrate the value of these four parameters (see Appendix A in S1 Appendix). The initial concentration of 𝑆 was fixed arbitrary at 𝑆_0_ = 1 cell/mL for naive animals and the ASCs generation rate was fixed at 𝛾 = 1 × 10^―5^ cells/virion/day [59]. The decay rate of ASCs and antibodies were fixed to 𝛿_𝑆_ = 0.31 day^―1^ and 𝛿_𝐴𝑏_ = 0.058 day^―1^ [49], corresponding to half-lives of 2.3 and 12 days, respectively. A profile likelihood was performed to validate the values of the three parameters 𝑆_0_, 𝛾 and 𝛿_𝑆_ (see Appendix C in S1 Appendix). All other parameters were estimated.

**Table 1.** Definition of model parameters TCID50 corresponding to a concentration of 2.80 × 10^8^ virions/ml measured just after inoculation in a 5mL sample.

| Parms | Description | Unit | Value | Refs |
| --- | --- | --- | --- | --- |
| **Viral dynamics model** |  |  |  |  |
| $\beta_0$ | Viral infectivity rate in absence of efficient immune response | mL/copies/day | Estimated | |
| $\delta$ | Death rate of infected cells | day<sup>-1</sup> | Estimated | |
| $P$ | Viral production rate | Virions/cell/day | Estimated | |
| $\alpha_{sgRNA}$ | Infected cells and sgRNA ratio | Virions/cell | Estimated | |
| $k$ | Eclipse rate | day<sup>-1</sup> | 3 Fixed* | [32] |
| $c$ | Clearance of produced virions | day<sup>-1</sup> | 3 Fixed* | [32] |
| $c_i$ | Clearance of inoculated virions | day<sup>-1</sup> | 20 Fixed* | [32] |
| $\mu$ | Percentage of infectious virions | Unitless | $10^{-3}$ Fixed | [32, 65] |
| $T_0^{N,nbc}$ | Initial number of target cells (Naso) | cell | $1.25 \times 10^5$ Fixed | [32, 41] |
| $T_0^{T,nbc}$ | Initial number of target cells (Trachea) | cell | $2.25 \times 10^4$ Fixed | [32, 41] |
| $Inoc_0$ | Number of inoculated virions** | virions | $1.40 \times 10^9$ Fixed | |
| **Humoral response model** |  |  |  |  |
| $\gamma$ | ASCs generation rate | Cells/virions/day | $1 \times 10^{-5}$ Fixed* | [59] |
| $\rho$ | Proliferation rate of ASCs | day<sup>-1</sup> | Estimated | |
| $\delta_{Ab}$ | Elimination rate of antibodies | day<sup>-1</sup> | 0.058 Fixed | [49, 84] |
| $\theta$ | Antibody production rate | AU/cell/day | Estimated | |
| $\alpha_{nab}$ | Proportion of nAb among bAb | unitless | Estimated | |
| $\eta$ | Net effect of nAb on virus neutralization | mL/AU | Estimated | |
| $\delta_S$ | Elimination rate of ASCs | day<sup>-1</sup> | 0.31 Fixed* | [49] |
| $S_{max}$ | Carrying capacity of ASCs | Cells/mL | Estimated | |
| $S_0$ | Initial ASCs concentration in Naïve NHPs | Cells/mL | 1 Fixed* | |

ASCs, antibody-secreting cells; AU, arbitrary unit; bAb, binding antibody; nAb, neutralizing antibody; NHP, non-human primate. \* Value calibrated by profile likelihood. \*\*Inoculation of $10^5$ TCID<sub>50</sub> corresponding to a concentration of $2.80 \times 10^8$ virions/ml measured just after inoculation in a 5mL sample.

### Observation model

To describe the log_10_-transformed dynamics of the viral genomic RNA, 𝑌^1^, and the subgenomic RNA, 𝑌^2^, in the two URT compartments, we kept the mathematical model previously proposed [32]. In particular, as shown in Equations (10) and (11), the gRNA was defined as the sum of inoculated (𝑉_𝑠_), infectious (𝑉_𝑖_) and non-infectious (𝑉_𝑛𝑖_) virions, while the sgRNA was described as a proxy of infected cells (𝐼_1_ + 𝐼_2_). For the humoral response, the model directly describes the log_10_ transformation of the concentrations of binding, 𝑌^3^, and neutralizing antibodies, 𝑌^4^, in AU/mL. Accordingly, for the 𝑖^𝑡ℎ^animal (𝑖 = 1 ,⋯,𝑁) at the 𝑗^𝑡ℎ^ time point (𝑗 = 1,⋯𝑛_𝑖_), the structural model is given as

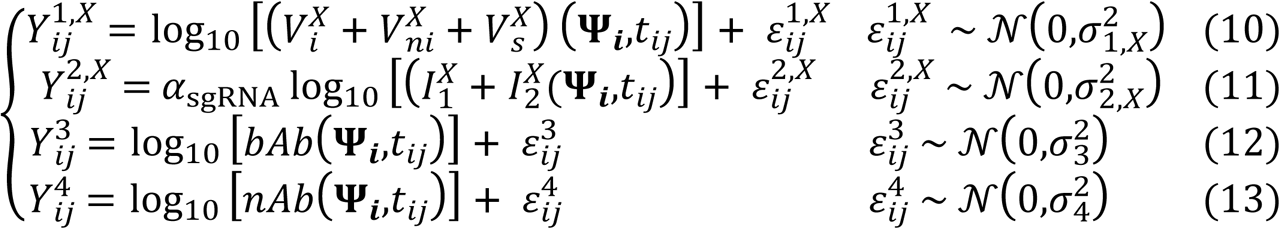

where 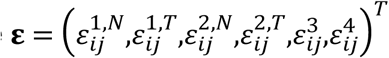 is the additive residual Gaussian error of mean 𝟎 and diagonal variance-covariance matrix 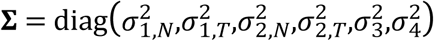 The vector 𝚿_𝐢_ corresponds to the individual value for the animal 𝑖 of the vector of model parameters 𝚿.

### Statistical model

#### General model

To account for variability observed on individual viral and antibody dynamics, some parameters of 𝚿 have to be defined as treatment group- and/or individual-specific. To this end, a statistical model is built on which model parameters are assumed to be log-transformed to ensure their positivity, except for 𝛼_𝑛𝑎𝑏_which follows a logit-normal distribution bounded between 0 and 1, and are described by a mixed-effects model. Accordingly, each estimated parameter 𝜓 of 𝚿, except for 𝛼_𝑛𝐴𝑏_, is defined as:

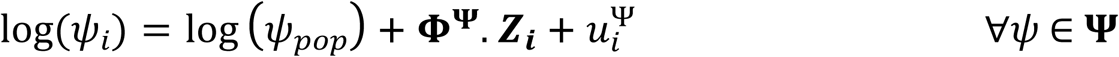

where 𝜓_𝑝𝑜𝑝_ is the fixed effect, i.e. the mean value in the population, 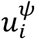 represents the random effect assumed normally distributed of mean zero and standard deviation 𝜔^𝜓^and independent from random effects of other parameters. The variable 𝐙_𝐢_ is the vector of group of NHP 𝑖, such that 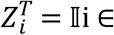 (Naive,CD40.RBDv ― Naive,Conv, CD40.RBDv ― Conv, CD40.PanCoV ― Conv,mRNA ― Conv ) , and 𝚽^𝛙^is the matrix of regression coefficients related to covariates for which the parameter 𝜓 has been adjusted for. Table 2 describes all categorical covariates that have been considered in the work.

**Table 2.**
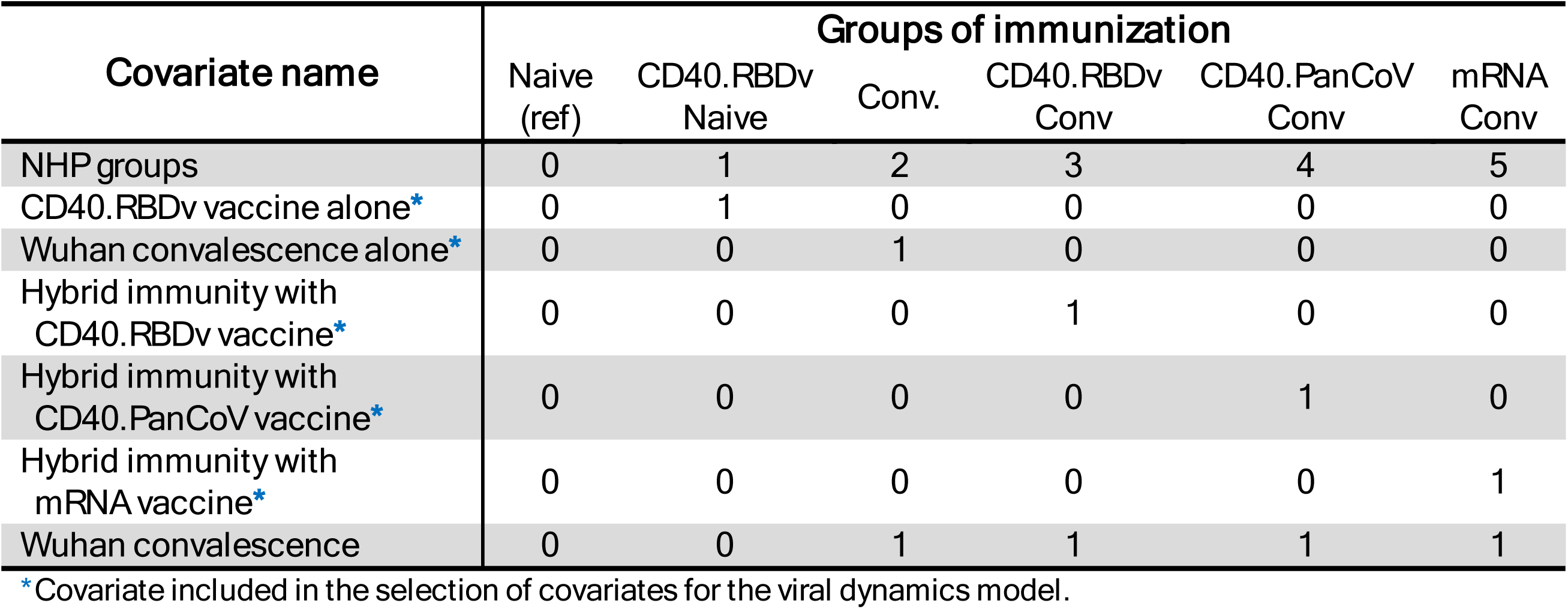
Definition of categorical covariates tested. Each line represents a covariate and columns represent categories built from the six groups of NHPs. The value 0 indicates that the group belongs to the group of reference

| Covariate name | Groups of immunization |  |  |  |  |  |
| --- | --- | --- | --- | --- | --- | --- |
|  | Naive (ref) | CD40.RBDv Naive | Conv. | CD40.RBDv Conv | CD40.PanCoV Conv | mRNA Conv |
| NHP groups | 0 | 1 | 2 | 3 | 4 | 5 |
| CD40.RBDv vaccine alone* | 0 | 1 | 0 | 0 | 0 | 0 |
| Wuhan convalescence alone* | 0 | 0 | 1 | 0 | 0 | 0 |
| Hybrid immunity with CD40.RBDv vaccine* | 0 | 0 | 0 | 1 | 0 | 0 |
| Hybrid immunity with CD40.PanCoV vaccine* | 0 | 0 | 0 | 0 | 1 | 0 |
| Hybrid immunity with mRNA vaccine* | 0 | 0 | 0 | 0 | 0 | 1 |
| Wuhan convalescence | 0 | 0 | 1 | 1 | 1 | 1 |

\*Covariate included in the selection of covariates for the viral dynamics model.

### Model building approach

Because of identifiability issues, random and covariates effects have not been added on all model parameters. The data-driven approach applied to build the optimal statistical model to fit viral and humoral data is described in Figure 3.

**Figure 3.**
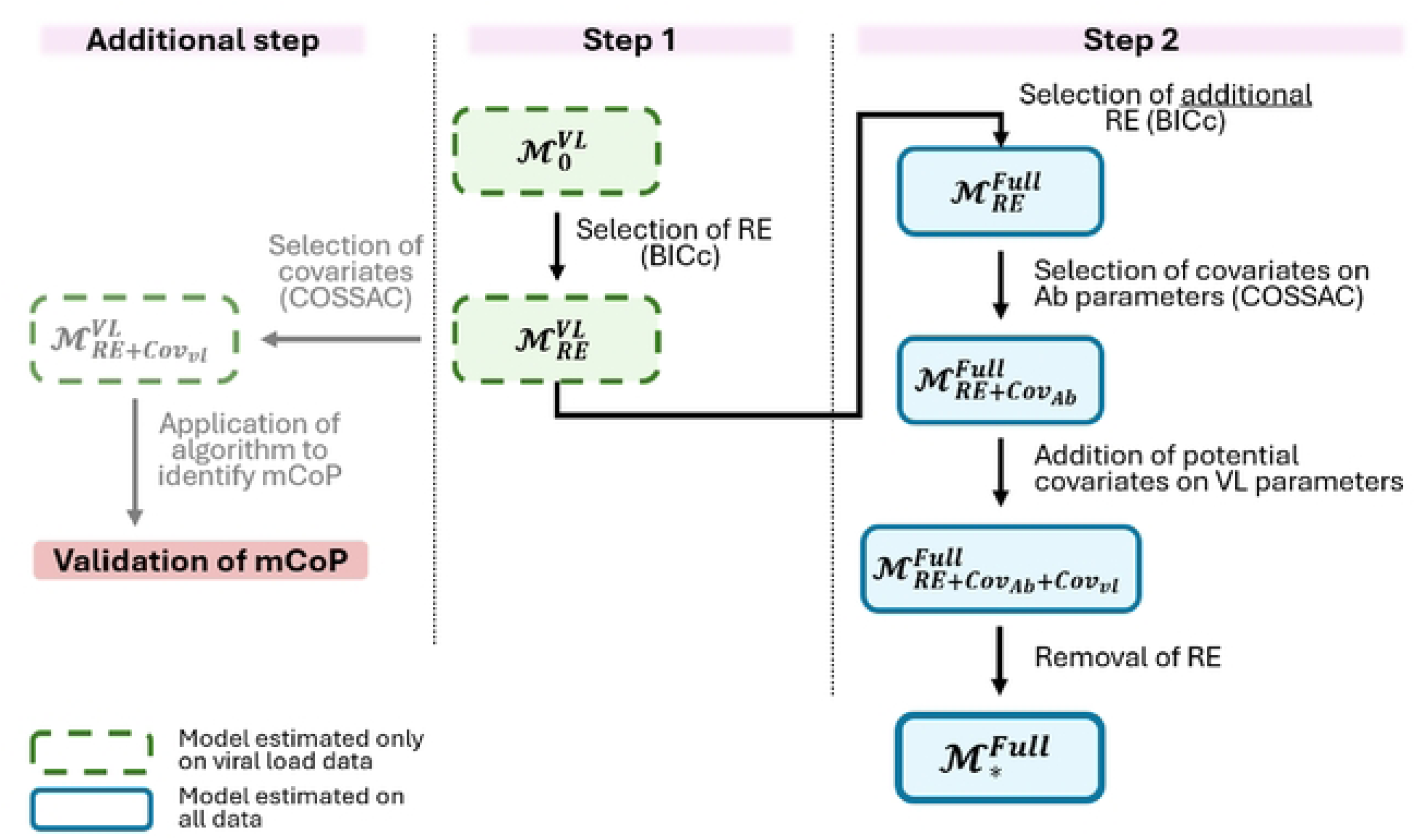
Two-step statistical model building approach applied. Rectangles represent models with green colored being based on the viral model (ℳ^VL^) described by equations (1)-(6) and blue colored the full model (ℳ^Full^) jointly describing viral and humoral dynamics by equations (1)-(9). Subscripts in model names indicate elements included in the statistical model: 0 for empty model, RE for random effects, Cov_vl_ for covariates on viral parameters, Cov_Ab_ for covariates on antibody parameters and ∗ for the final optimal model. Notations: *BICc*, corrected Bayesian Information Criterion; *COSSAC*, Conditional Sampling use for Stepwise Approach on Correlation tests; *mCoP*, mechanistic correlate of protection.

The first step consisted in building the statistical model of the viral dynamics model, i.e., focusing on 𝚿_𝐯𝐥_ and using only viral dynamics data (see Appendix A in S1 Appendix for a full description). Firstly, random effects were selected by estimating all potential models and using the corrected Bayesian information criterion (BICc) [66] as selection criterion (the lower the better). Secondly, immunological background-specific effects on viral parameters were selected by applying the Conditional Sampling use for Stepwise Approach on Correlation tests (COSSAC) algorithm [67]. In particular, this selection validated the blockage of new infections and the enhancement of infected cells elimination as the main immune mechanisms allowing the control of viral infection. Furthermore, the application of the methodology we developed to identify mechanistic CoP [32] confirmed the implication of antibody responses in the capture of this first mechanism (see Appendix B in S1 Appendix). Once random effects selected on viral parameters, the second step consisted in including the antigen-mediated immune response in the statistical model considering the full model and using viral and antibody data (see Appendix C in S1 Appendix for a full description). We first selected random effects on antibody dynamics parameters, i.e., on 𝚿_𝐴𝑏_, by applying a forward selection algorithm using BICc as selection criterion. Then, effects of vaccination and/or Wuhan primo-infection on the humoral responses were identified by applying the COSSAC covariate selection algorithm. In this step, no adjustment for covariates on viral dynamics parameters were allowed. Moreover, as aforementioned, we forced the adjustment of (𝑆_0_) for the groups of immunization. Once covariates selected, the significance of effects were verified by Wald test non-significant ones were removed. Finally, the addition of covariates on viral parameters was then tested before removing unnecessary random effects using a backward selection approach with BICc criterion.

#### Selected statistical model

The optimal statistical model identified accounted for inter-individual variability on the parameters 𝛿, 𝑆_0_, 𝜃 and 𝜂, and covariate effects on parameters 𝛿, 𝑆_0_, 𝜌, 𝜃 and 𝜂. In particular, in addition to the group effects on 𝑆_0_, we identified 1) an effect of Wuhan infection on the proliferation rate of ASCs 𝜌, 2) an effect of hybrid immunity on the neutralization effect 𝜂, 3) an effect of the groups Convalescent and CD40.RBDv-Naive on the antibody production rate 𝜃, and 4) an effect of the Convalescent group on the elimination rate of infected cells 𝛿. Accordingly, the final statistical model is given by the following equations:

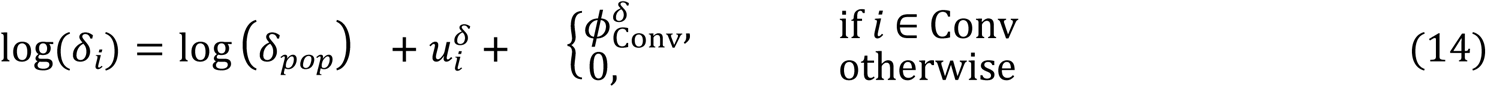

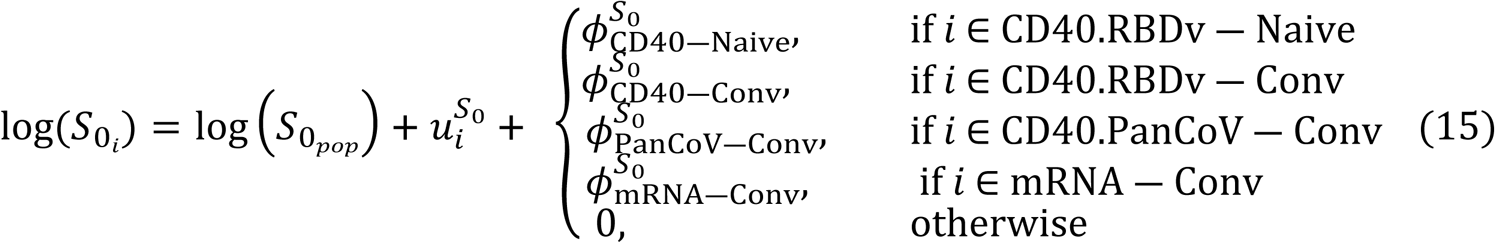

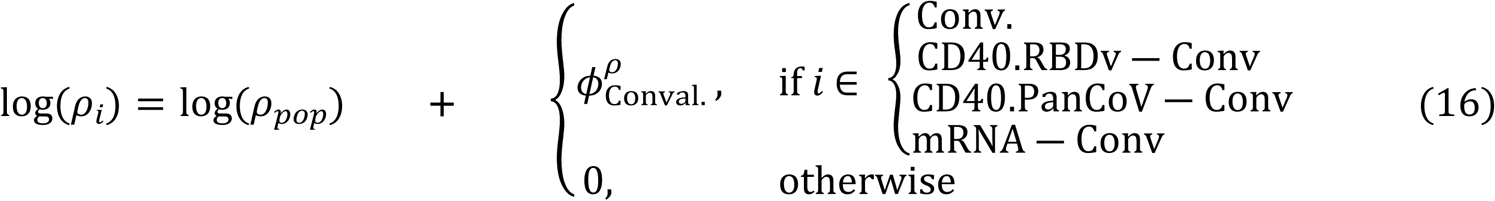

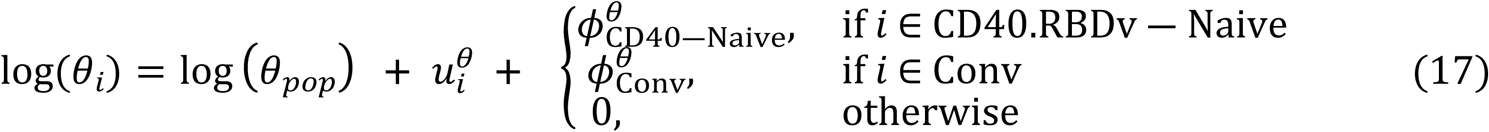

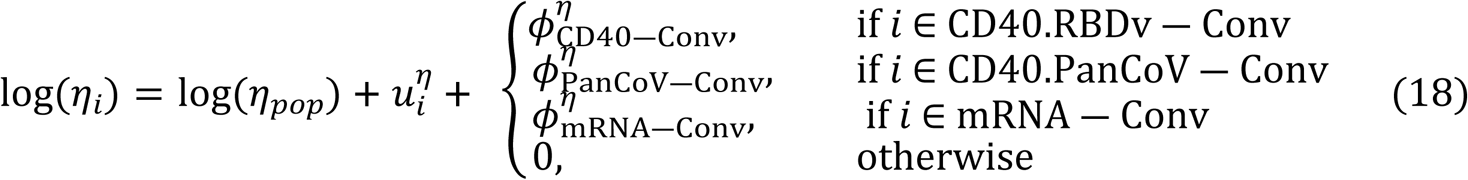

## Model identifiability

Before the estimation of the full model, we investigated its structural identifiability. We first analyzed the global structural identifiability of the full model given by Equations (1) to (9). However, we encountered out-of memory issues induced by the large number of state variables and the small number of observables. To overcome this issue, we first analyzed the global identifiability of the reduced version of the model involving only a single URT compartment. Second, we verified the local identifiability of the full model. All analyzes were performed considering the parameters 𝜇, 𝑘, 𝑐 and 𝑐_𝑖_ as fixed. The analysis of the reduced model highlighted the non-identifiability of the set of parameters {𝛿_𝑆_,𝛾,𝜌,𝑆_𝑚𝑎𝑥_,𝜃}, and identified the three following combinations of parameters as identifiable: 𝛾𝜃, 𝛿_𝑆_ ― 𝜌, 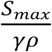. Accordingly, we chose to fix 𝛿_S_ and 𝛾 making the model globally identifiable. Finally, we validated the local identifiability of the full model under these conditions.

### Binding inhibitory functionality allowing protection

#### Derivation of the protective threshold

Through this work, we aimed at deriving a protective threshold of the immune marker previously identified as a consistent mechanistic correlate of protection against SARS-CoV-2 infection. In other words, we want to quantify the minimal concentration of antibodies inhibiting ACE2/RBD binding required to rapidly control viral replication after exposure and thus avoiding the establishment of viral infection within an animal. To determine this protective threshold, noted 𝜏_𝑛𝐴𝑏_, we used the basic within-host reproduction number ℛ_0_ that represents the number of infected cells newly generated from one infected cell introduced in a full population of susceptible cells (i.e., uninfected target cells here) at the time of infection, hence 𝑇 = 𝑇_0_. To derive the formula of the reproduction number, we applied the next-generation method [68,69] in which ℛ corresponds to the spectral radius, i.e., the dominant eigen value, of the next-generation matrix. By independence of the infection system within the two URT compartments, we can demonstrate that each compartment can be studied separately allowing the derivation of two basic reproduction numbers, 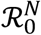 and 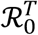 for nasopharyngeal and tracheal viral infections, respectively. The global ℛ_0_ is then given by 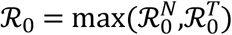. By applying the next-generation method (see Appendix D in S1 Appendix for a full description), the reproduction number in the compartment 𝑋 is given by

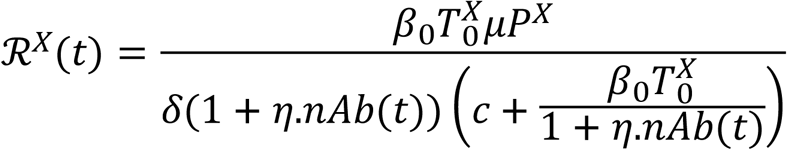

which depends on the dynamics of neutralizing antibodies and thus on time [70]. Accordingly, the global basic reproduction number is given by

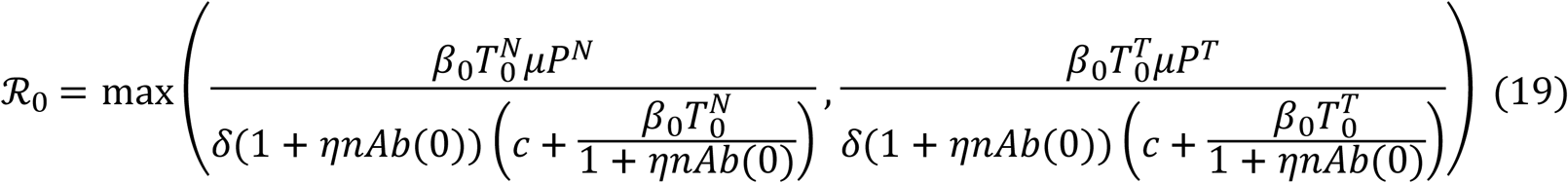

The protective threshold 𝜏_𝑛𝐴𝑏_represents the smallest concentration of neutralizing antibodies required at the time of infection allowing the control of viral infection meaning such that ℛ_0_ ≤ 1. The reproduction number being a decreasing function of 𝑛𝐴𝑏, this threshold corresponds to the value of 𝑛𝐴𝑏(0) requested to reach ℛ_0_ = 1, giving the following.

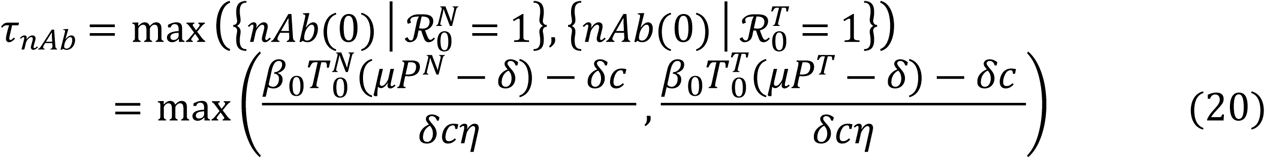

## Simulation of the impact of the immunological background on the control of viral replication and the protective threshold

Next, we used model parameter estimates to analyze the ability of NHPs to control viral replication after Delta infection. In particular, we provided the distribution, i.e. mean value and confidence interval, for both the basic reproduction number ℛ_0_and the protective ACE2/RBD binding inhibition threshold 𝜏_𝑛𝐴𝑏_in each NHP group. To derive confidence intervals accounting for uncertainty in population parameter estimation, we sampled 𝐾 = 500 population parameters from their posterior distribution. Then, for each set of simulated parameters (i.e., without random effects, 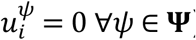), we calculated ℛ and 𝜏 using Equations (19) and (20), respectively, with 𝑛𝐴𝑏(0) given by 𝛼_𝑛𝑎𝑏_𝜃𝑆_0_/𝛿_𝐴𝑏_in ℛ_0_. Subsequently, we derived the mean protective threshold and its 95% confidence interval requested in each NHP group at the time of Delta infection to control viral replication.

## Simulation of counterfactual scenarios

Describing the dynamics of biological processes, mechanistic models have the major advantage after their estimation to be used to explore counterfactual scenarios. Accordingly, we used the model to simulate hypothetical scenarios for 1) analyzing and interpreting the distinct value of protective threshold estimated in each NHP group, and 2) understanding which immune mechanisms drive viral control regarding the immunological background. To account for uncertainty in population parameters estimation in simulations, we sampled 𝐾 = 500 sets of population parameters using their posterior distribution. For each set of parameters, we predicted the counterfactual genomic and subgenomic viral load trajectories and calculated the aforementioned viral load dynamics descriptors (i.e., values and time to peak VL, AUC, duration of clearance and acute stages). The mean and 95% confidence interval of trajectories, and the median and inter-quartile range of genomic and subgenomic VL descriptors were calculated in each NHP group. All scenarios were evaluated with the same sets of sampled population parameters and compared to the factual scenario. As described in Table 3, three distinct counterfactual statements have been studied in this work to analyze viral control properties: "What would have been viral control if":

1. "similar levels of neutralizing antibodies had been observed in all NHP groups at exposure."
2. "no uninfected target-cell depletion occurred after infection."
3. "neutralizing antibodies were inefficient in neutralizing viruses."

**Table 3.** Description of the counterfactual scenario. Distinct counterfactual scenarios have been analyzed to understand what would have been viral control in NHP groups under different conditions.

| Scenario | *What if ...* Statement | Modifications applied to all NHP groups |
| --- | --- | --- |
| | What if **similar levels of nAb** (and by extension similar bAb) had been observed at exposure in all NHP groups ?<br>Three distinct levels of antibodies have been tested: | $nAb_0 = nAb_{\text{fixed}}$ |
| **LownAb** | ▷ **Low level of nAb** with $nAb_{\text{fixed}}$ equals to the level of the protective threshold found in CD40.PanCoV-Conv group to protect at least 95% of NHPs. | $nAb_{\text{fixed}} = 0.90 \text{ AU/mL}$ |
| **MidnAb** | ▷ **Medium level of nAb** with $nAb_{\text{fixed}}$ equals to the median level of nAb observed in CD40.PanCoV-Conv group at exposure. | $nAb_{\text{fixed}} = 24 \text{ AU/mL}$ |
| **HinAb** | ▷ **High level of nAb** with $nAb_{\text{fixed}}$ equals to the maximum level of nAb observed in CD40.PanCoV-Conv group at exposure. | $nAb_{\text{fixed}} = 437 \text{ AU/mL}$ |
| **Nodepl** | What if **no depletion of uninfected target-cells** occurred after viral Delta infection ? Accordingly, viral control is assumed to be only driven by antibody neutralizing activity. | $T(t) = T_0 \quad \forall t$ |
| **Noneut** | What if **nAb are inefficient** in their activity to **neutralize viruses** and consequently to reduce viral infectivity? | $\eta = 0 \Rightarrow \varepsilon_{Ab}(t) = 0 \quad \forall t$ |

## Implementation procedure

All model parameters were estimated with the stochastic approximation expectation-maximization (SAEM) algorithm implemented in Monolix software version 2023R1 [71,72]. Likelihood was estimated by importance sampling and standard errors were obtained by stochastic approximation of the Fisher Information Matrix using a Markov chain Monte Carlo algorithm. Statistical significance of covariates included in the statistical model were evaluated by a Wald test. All analyses were performed using R version 4.2.1 [73] and Monolix estimations were performed through the lixoftConnectors R package [74]. Simulations of counterfactual scenarios were performed using Simulx software version 2023R1 [75] via the lixoftConnectors [74] and RsSimulx [76] R packages.

Structural identifiability of the model was performed using Julia version 1.11.3 [77] and the library *StructuralIdentifiability* [78].

## Results

### Viral and antibody kinetics according to immunological background

In our preclinical study, genomic and subgenomic RNA were quantified at regular time points during the 30 days following SARS-CoV-2 Delta challenge. After exposure, all animals developed rapid infection with genomic RNA peaking around 1 and 3 d.p.e in both the nasopharynx (𝑝 = 0.197) and trachea (𝑝 = 0.263) (Figures 4A-B, Figure 5A and Figure S4A). Afterwards, viral loads in the two URT compartments declined until reaching undetectable levels. At peak, viral load levels were higher in naive animals either vaccinated (CD40.RBDv-Naive) or not and convalescent animals than in vaccinated NHPs (CD40.RBDv-Conv, CD40.PanCoV-Conv, mRNA-Conv) with hybrid immunity. The same profiles were observed in the trachea (Median, log_10_ copies/mL: 8.28 in Naive, 7.52 in CD40.RBDv-Naive, 6.62 in Convalescent, 5.09 in CD40.RBDv-Conv, 4.08 in CD40.PanCoV-Conv and 3.82 in mRNA-Conv, 𝑝 < 0.001) or in the nasopharynx (Median, log_10_ copies/mL: 8.50 in Naive, 7.87 in CD40.RBDv-Naive, 7.38 in Convalescent, 4.03 in CD40.RBDv-Conv, 3.52 in CD40.PanCoV-Conv and 3.18 in mRNA-Conv, 𝑝 < 0.001). Similarly, shorter duration of clearance and acute stages were observed in the case of hybrid immunity with, for instance, their median durations ranging between 1-2 days in the nasopharynx while mean clearance and acute stages lasted 9-17 days and 6-16 days in the three other groups, respectively (Figure 5A).

**Figure 4.**
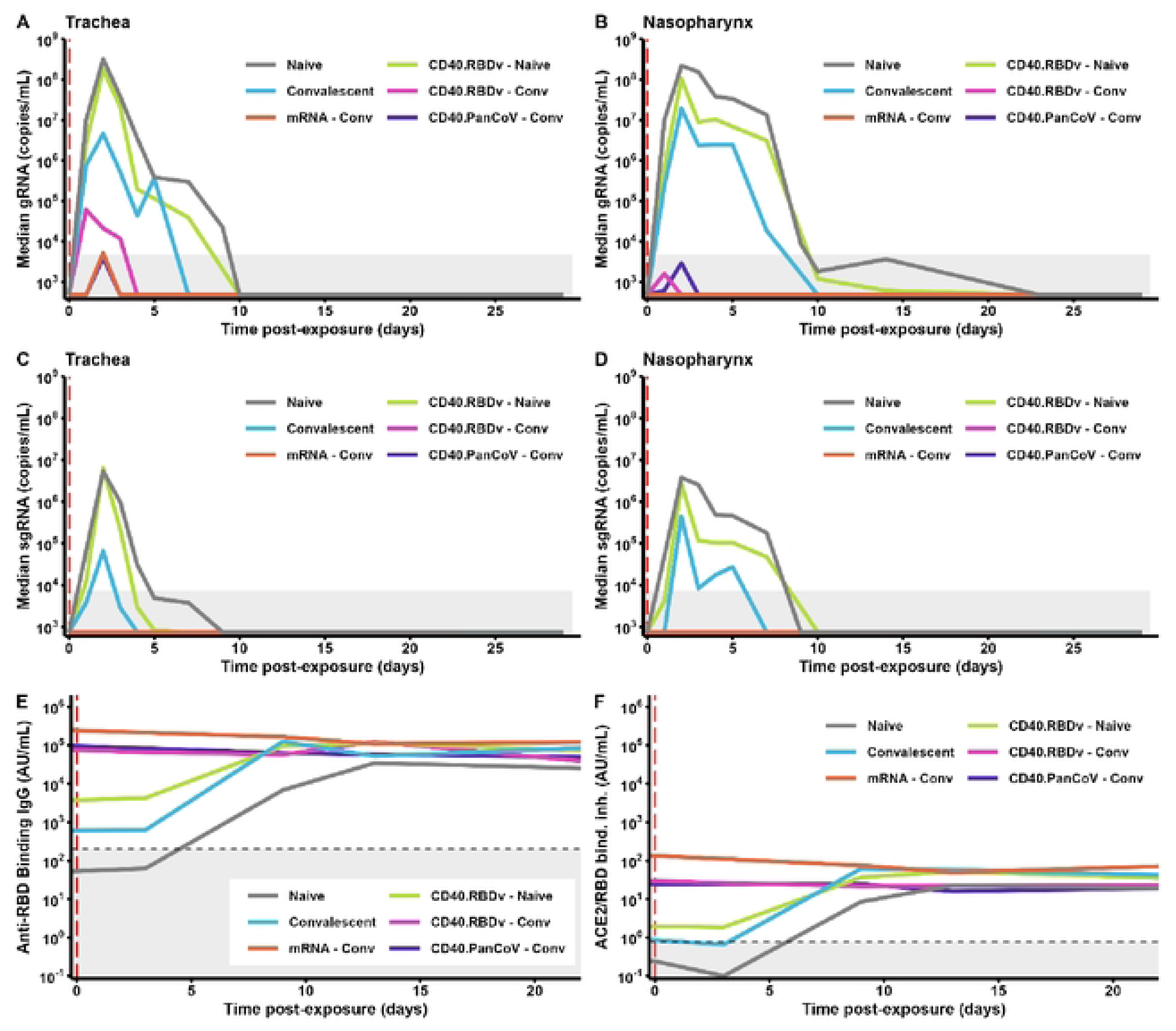
Median viral and antibody dynamics observed after SARS-CoV-2 infection. (A) Dynamics of gRNA viral load observed in trachea (in copies/mL). (B) Dynamics of gRNA viral load in nasopharynx (in copies/mL). (C) Dynamics of sgRNA viral load observed in trachea (in copies/mL). (D) Dynamics of sgRNA viral load in nasopharynx (in copies/mL). (E) Dynamics of anti-RBD binding antibodies (in AU/mL). (F) Dynamics of antibodies neutralizing ACE2/RBB binding (in AU/mL). In all subplots, median dynamics (in log_10_ scale) are observed in Naive (gray), CD40.RBDv - Naive (light green), Convalescent (blue), CD40.RBDv - Conv (pink), CD40.PanCoV - Conv (purple) and mRNA - Conv (orange) animals. Vertical red dashed highlights the time of exposure, and shaded areas highlight the limits of quantification for viral loads (A-D) and limits of positivity for antibodies (E-F).

**Figure 5.**
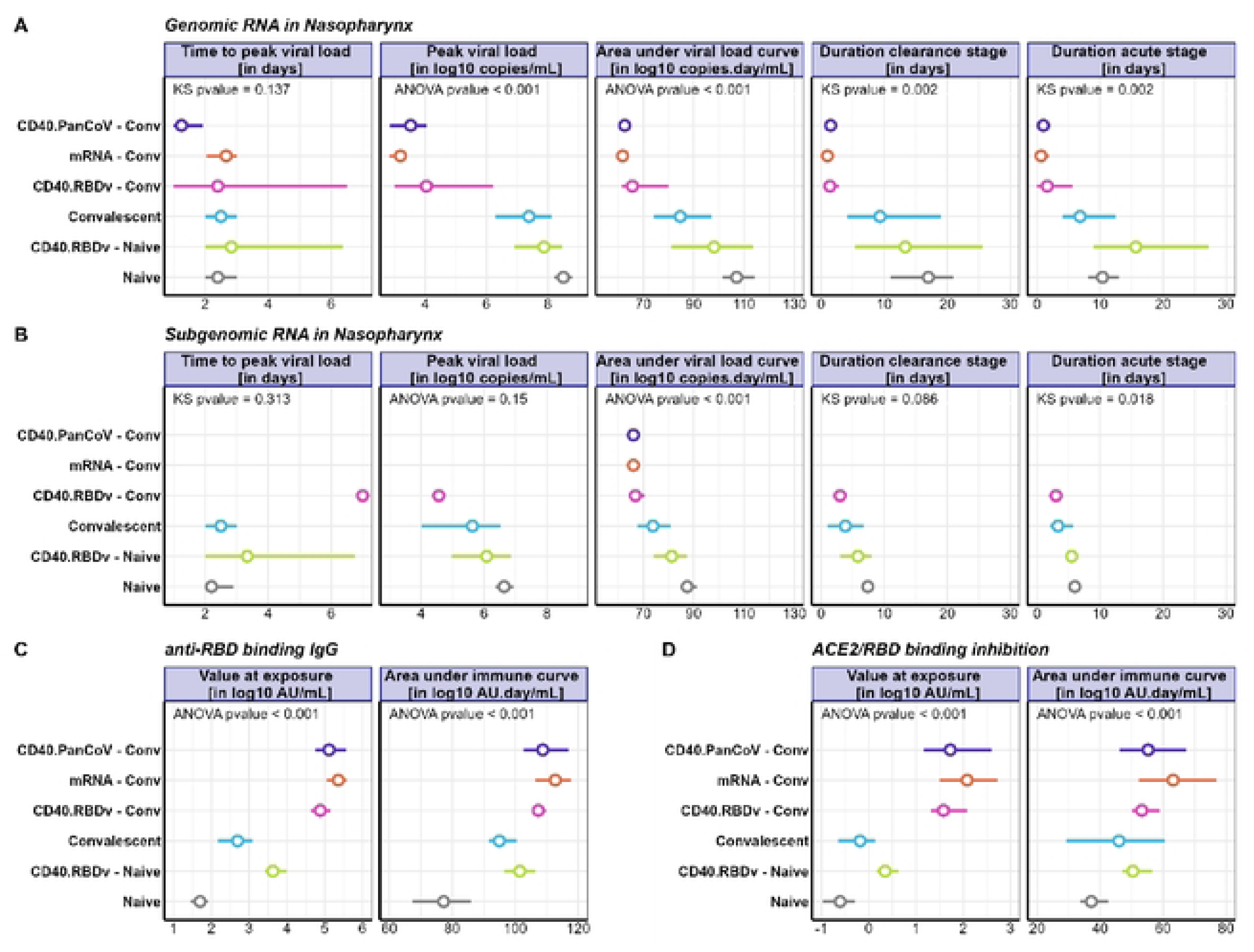
Comparison of viral and antibody dynamics descriptors between NHP groups. (A) Descriptors of genomic RNA viral load dynamics in nasopharynx. (B) Descriptors of subgenomic RNA viral load dynamics in nasopharynx. (C) Descriptors of anti-RBD binding antibody dynamics. **(D)** Descriptors of ACE2/RBD binding inhibition dynamics. (A-B) Viral dynamics are described through 5 descriptors: time (in day) and value (in log_10_ cp/mL) of viral load peak, AUC of viral load curve (in log_10_ cp.day/mL), duration of clearance and acute stages (in day). (C-D) Antibody dynamics are described through 2 descriptors: value observed at exposure (in log_10_ AU/mL) and AUC of immune curve (in log_10_ AU.day/mL). (A-D) Circles and solid lines represent mean values and 95% confidence intervals, respectively. Adjusted p-values for either global ANOVA or Kruskal-Wallis comparison tests are provided for each descriptor.

Regarding subgenomic RNA dynamics characterizing viral replication following infection, undetectable viral loads were observed in the three hybrid groups (Figures 4C-D) while sgRNA viral loads peaked around 2 d.p.e in other groups, before decreasing and reaching undetectable levels after median times of 1.7 and 3.8 d.p.e in Convalescent, 4.3 and 5.8 d.p.e in CD40.RBDv-Naive, and 7.0 and 7.4 d.p.e in Naive NHPs, in the trachea and the nasopharynx, respectively (Figure 5B and Figure S4B). Thus, evaluation of sgRNA dynamics following Delta challenge highlighted a better control of viral replication in the context of hybrid immunity than in naive immunity.

In addition to viral loads, both anti-RBD binding IgG and ACE2/RBD binding inhibition were longitudinally quantified following Delta infection to evaluate antibody responses. As shown in Figures 4E-F, similar dynamics of the two immune markers were observed in the three groups of hybrid immunization, with mean baseline values ranging between 5 and 5.4 log_10_ AU/mL for binding antibodies, and between 1.7 and 2.3 log_10_ AU/mL for neutralizing antibodies (Figures 4E-F and Figures 5C-D). Antibody levels were significantly lower at exposure in the three other groups with mean values ranging from 1.7 log_10_AU/mL in Naive NHPs to 3.7 log_10_AU/mL in CD40.RBDv-Naive NHPs for binding IgG (𝑝 < 0.001), and from -0.54 log_10_AU/mL to 0.38 log_10_ AU/mL for neutralizing IgG (𝑝 < 0.001). Accordingly, 100% and 83% of Naive and Convalescent animals showed binding antibodies below the limit of positivity at exposure, respectively, and 100% and 50% for neutralizing antibodies (Table S1). These low antibody levels were followed by rapid increases until reaching similar concentrations in immunized animals after 10 to 14 d.p.e (Figures 4E-F).

## Mechanistic modeling of viral and humoral responses

### Preliminary result - Validation of antigen-induced immune mechanisms and mCoP

We first evaluated the viral model in absence of antigen-mediated immune response to identify immune mechanisms impacted by vaccine- and Wuhan convalescence-induced immunities (see Appendix A in S1 Appendix). In accordance with our previous work [32], we identified the blockage of new cells infection and the enhancement of infected cells elimination as the main mechanisms. In particular, the hybrid immunity allowed to significantly reduce viral infectivity 𝛽 with values divided by factors ranging from 2 000 to 60 000 in the three convalescent NHP groups compared to naive animals. We also identified an increase of the clearance of infected cells 𝛿 in Convalescent group of 115% compared to Naive group. The reader can refer to Appendix A in S1 Appendix for a full description of viral model estimation.

The application of our methodology to identify mCoP allowed us to validate our previous results [32] (see Appendix B in S1 Appendix for a full description of the applied methodology and results). We first showed the ability of antibody responses to capture the effects of hybrid immunity on the viral infectivity rate. Then, we demonstrated that antibody responses (both binding and neutralizing antibody dynamics) fulfilled criteria defining mCoP to explain the blockage of new target infection after immunization, whether by infection or vaccination. Thereafter, we provided evidence that the effect of previous immunity in Convalescent found on the elimination rate of infected cells 𝛿 remained unexplained after the adjustment of the model for antibody responses. Finally, we pointed out the benefit of considering immunity as a dynamical process by integrating the dynamics of immune markers to fully capture the effect of antibody responses on the viral dynamics. Accordingly, these results validate the necessity of joint modeling of viral and antibody dynamics to rightly quantify a protective threshold of the mCoP allowing rapid control of viral replication following Delta SARS-CoV-2 infection.

## Joint mechanistic modeling

The mathematical model we proposed to evaluate the impact of the immune response on the viral dynamics to quantify the CoP was simultaneously fitted to viral and humoral data collected in the 34 NHPs. Estimation of model parameters are gathered in Table 4. In absence of efficient immune responses (i.e. 𝑛𝐴𝑏 = 0 AU/mL), we estimated the viral infectivity 𝛽_0_ at 7.91 × 10^―5^ (CI_95%_ [6.27; 9.98] × 10^―5^) mL/copies/day, which is in the range of previously reported modeling results for naive animals [32,43,44,65]. The establishment of a robust humoral response allowed to significantly reduce the effective viral infectivity 𝛽 over time until reaching a mean value of 4.95 × 10^―6^ (CI_95%_ [1.41; 14.5

**Table 4.**
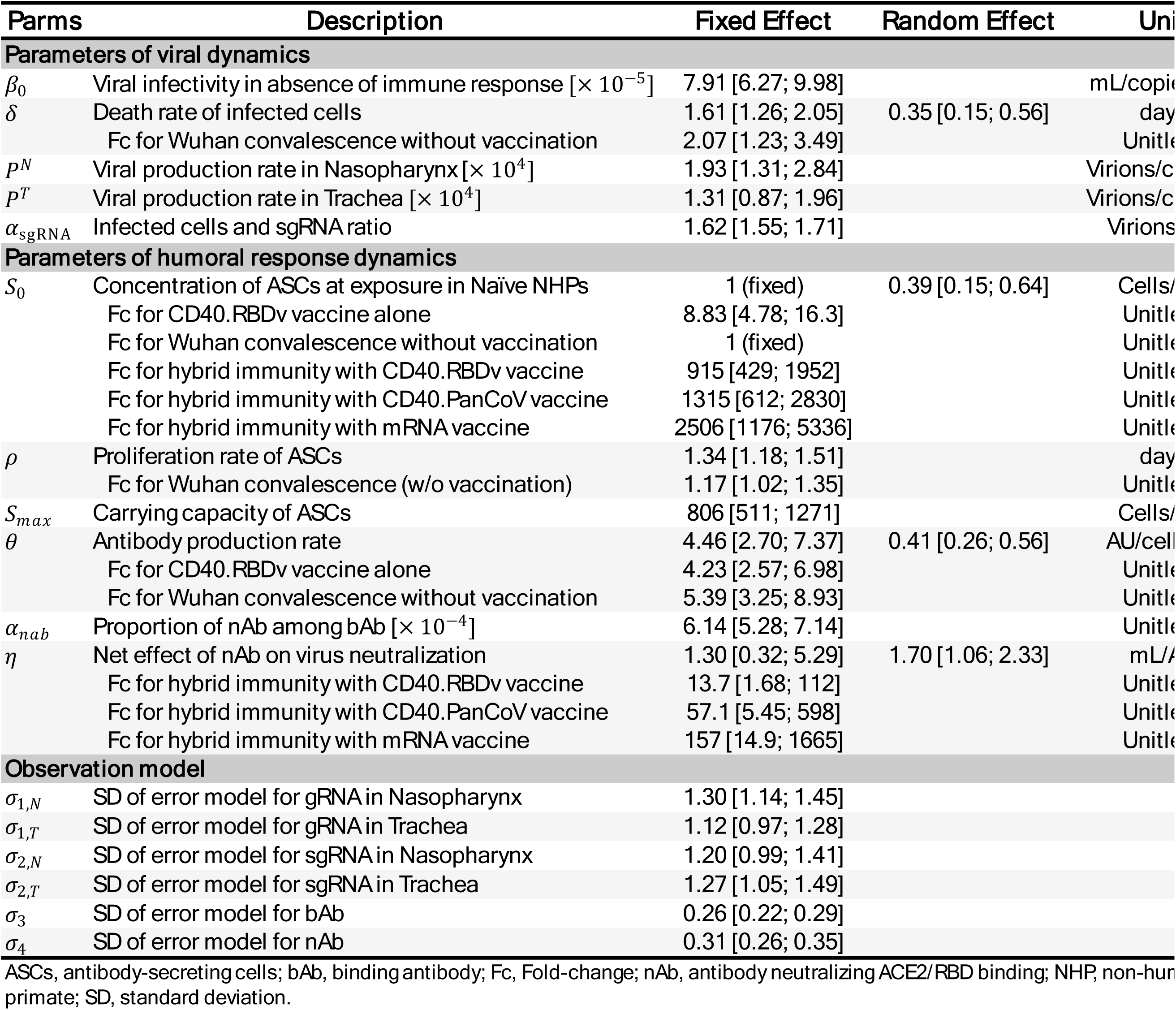
. Model parameter estimates.

] × 10^―6^) mL/copies/day in naive animals at 23 d.p.e (Figure S5A). An estimate of the elimination rate of infected cells 𝛿 of 1.61 (CI_95%_ [1.26; 2.05]) day^―1^ was found, which remains in accordance with the literature for SARS-CoV-2 with values ranging from 0.6 to 2 day^―1^ [32,41,43,44,65], and corresponds to a half-life of 10.3 (CI_95%_ [8.11; 13.2]) hours. Furthermore, we retrieved the effect induced by the Convalescent group on the elimination rate increasing 𝛿 by 107 % on average (𝑝 = 0.006). As initially identified in the viral model, a higher viral production rate was estimated in the nasopharynx (1.93 × 10^4^ (CI_95%_ [1.31; 2.84] × 10^4^) virions/cell/day) compared to the trachea (1.31 × 10^4^ (CI_95%_ [0.87; 1.96] × 10^4^) virions/cell.day) with a mean ratio of 1.47.

Regarding the humoral immune responses, several immunological background-specific covariates were identified on immune model parameters explaining the difference of antibody dynamics observed between NHP groups. First, we evaluated the effect of each group of immunization, compared to Naive animals, on the size of ASCs pool at exposure (S_0_). In particular, these effects allowed to capture the difference of antibody concentrations observed at baseline (before challenge) between animals. These initial concentrations of cells were significantly higher in all immunized groups (𝑝 < 0.0001), except for the Convalescent group, with values multiplied by a factor ranging from 8.83 (CI_95%_ [4.78; 16.32]) for CD40.RBDv-Naive group to 2506 (CI_95%_ [1176; 5536]) for mRNA-Conv group. The absence of significant differences in the Convalescent group (𝑝 = 0.67) is explained by the similarity of antibody concentrations observed at exposure between Naive and Convalescent animals (𝑝 = 0.033 for bAb0 and 𝑝 = 0.13 for nAb0, in natural scale; Figure 4 and Figure 5). Wuhan convalescence had an effect on the proliferation rate of ASCs, 𝜌, (𝑝 = 0.027) with a value increased by a factor 1.17 (CI_95%_ [1.02; 1.35]). Hence, regardless of the initial level of ASCs at exposure, the immunity established after the primary infection is more efficient to react against new infection by producing ASCs faster (Figure S5B). A higher production rate of antibodies by ASCs, 𝜃, was identified for NHPs belonging to CD40.RBDv-Naive and Convalescent groups compared to Naive animals. The production rate was increased by a mean factor of 4.23 (CI_95%_ [2.57; 6.98], 𝑝 < 0.0001) in vaccinated animals, and by 5.39 (CI_95%_ [3.25; 8.93], 𝑝 < 0.0001) in convalescent animals, resulting in a faster increase of antibody dynamics following viral infection. The absence of identification of this effect for animals with hybrid immunity could result from identifiability issues induced by the lack of observation of plasma cells and the simplicity of the model describing the saturation level of antibodies already reached at exposure (Figure 4).

## Impact of the immunological background on the efficacy of antibodies

The consideration of a joint mechanistic model integrating interactions between viral and humoral immune responses allowed us to identify an additional effect of immunological backgrounds on the functionality of antibodies to neutralize viruses. Indeed, the MSD V-Plex ACE2 Neutralization kit used to quantify neutralizing antibodies in this study, being only a quantitative test for a given variant, cannot be directly informative on antibody affinity or avidity. However, the simultaneous fitting of viral and nAb data allowed to inform on this nAb efficacy and to highlight a difference of efficacy between animals. Although no difference was found between Naive, Convalescent and CD40.RBDv-Naive animals, we distinguished significantly higher net effect of nAb on virus neutralization in animals with hybrid immunity, with values of 𝜂 increased by a factor 13.7 (CI_95%_ [1.68; 112], 𝑝 = 0.014) in CD40.RBDv-Conv, 57.1 (CI_95%_ [5.45; 598], 𝑝 < 0.001) in CD40.PanCoV-Conv, and 157 (CI_95%_ [14.9; 1665], 𝑝 < 0.0001) in mRNA-Conv groups. These effects first translate into a difference in neutralization efficacy over time between groups (Figure 6), characterized by the function 𝜀_𝐴𝑏_. A mean efficacy ranging between 5% and 63% at exposure in animals without hybrid immunity was observed before increasing over time until reaching 94 to 99% at 30 d.p.e. This increase results from the increasing dynamics of neutralizing antibodies observed after infection. At the opposite, stable efficacy levels closed to 100% were already found in the three other groups from the time of exposure. These high levels of neutralization efficacy explained the ability of immune system to more rapidly control viral replication in animals with hybrid immunity than others. Although this better neutralization efficacy could be explained by higher levels of antibodies measured in these three groups, the model also pointed out a better efficacy of antibodies to neutralize viruses after primo-infection and vaccination than in naive animals. Indeed, as shown in Figure 6B, a given level of nAb led to a higher percent of neutralization in case of hybrid immunity. For instance, the model predicted that a concentration of 0.1 AU/mL of nAb was able to neutralize only 11% (CI_95%_ [3%; 26%]) of viruses in Naive, CD40.RBDv-Naive and Convalescent animals, while the same level allowed to neutralize 57% (CI_95%_ [20%; 89%]), 83% (CI_95%_ [48%; 99%]) and 92% (CI_95%_ [72%; 99%]) of viruses in CD40.RBDv-Conv, CD40.PanCoV-Conv and mRNA-Conv NHPs. The maturation of the memory B-cell response enhanced by the primo-infection followed by vaccination could explain this better efficacy of neutralizing antibodies. Moreover, mRNA and CD40.PanCoV vaccines appeared as more efficient to stimulate memory responses after Wuhan convalescence than CD40.RBDv vaccine. Furthermore, these results highlighted the lack of improvement of the quality of humoral immune response after infection and or vaccination only in a naïve immune system as compared to hybrid immunity.

**Figure 6.**
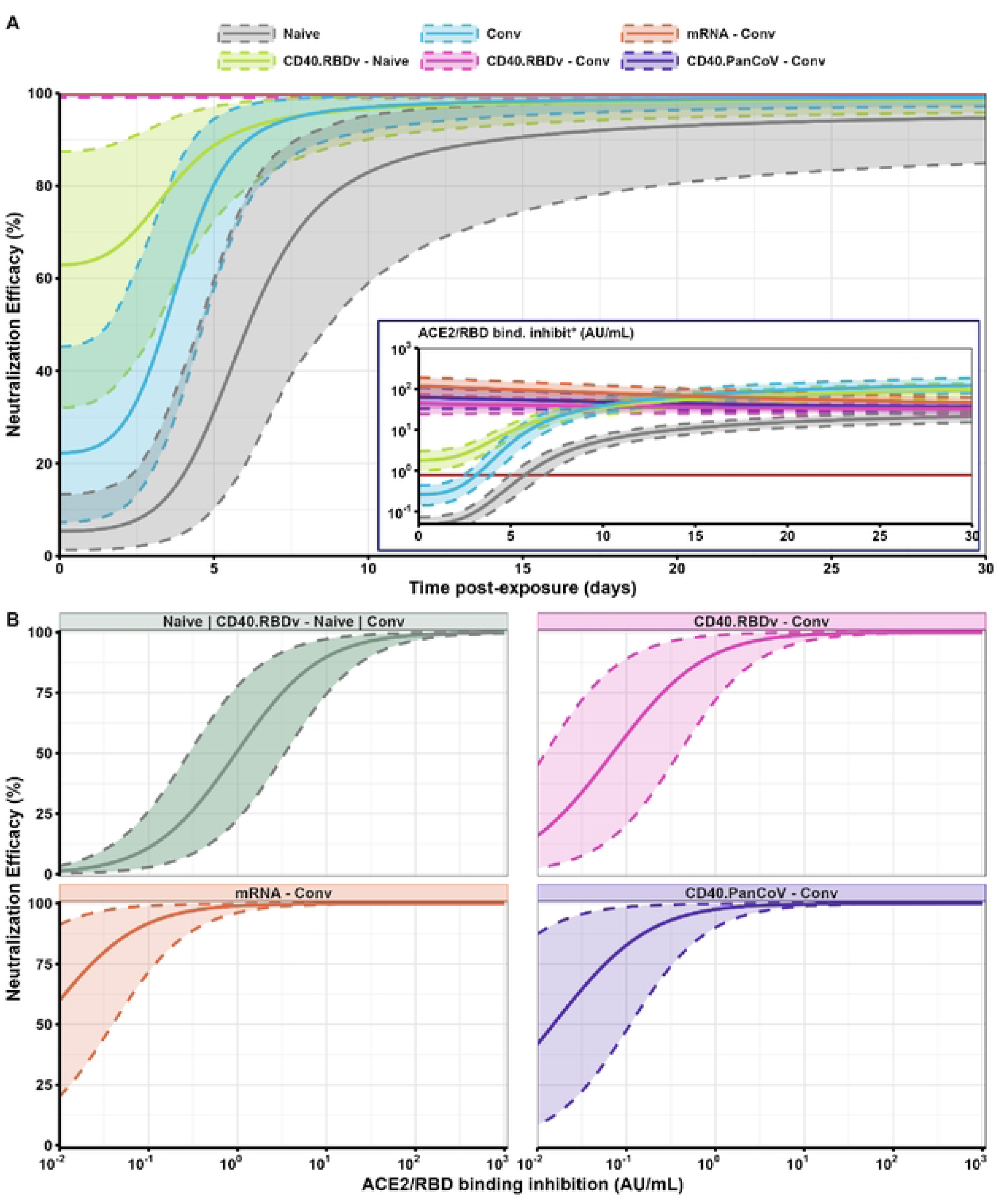
Predicted evolution of the neutralization efficacy of antibodies (in. %), 𝛆_𝐀𝐛_ = ℱ (𝐭,𝐧𝐀𝐛)**. (A)** Evolution of neutralization efficacy over time following SARS-CoV-2 infection. Dynamics of ε_Ab_ = f(nAb(t)) predicted in each NHP group, with embedded figure highlighting the related predicted dynamics of neutralizing antibodies, nAb(t), in AU/mL. **(B) Evolution of the neutralization efficacy of nAb according to the level of nAb.** Dynamics of ε_Ab_ = f(nAb) predicted in each NHP group for nAb values ranging from 10^―2^ to 10^3^ AU/mL. (A-B) Continuous thick lines and colored shaded areas represent mean predictions and 95% prediction intervals, respectively, for Naive (in gray), CD40.RBDv-Naive (in light green), Convalescent (in blue), CD40.RBDv-Conv (in pink), CD40.PanCoV-Conv (in purple) and mRNA-Conv (in orange) NHP groups.

## Benefit of modeling antibody response

The inclusion of antibody dynamics in the full mechanistic model and by allowing parameters of humoral response to differ between animals, with random effects identified on 𝑆_0_, 𝜃 and 𝜂, led to a fully capture of the individual variation of viral infectivity initially identified in the viral model (Appendix A in S1 Appendix). Indeed, no covariates or random effects were found on 𝛽_0_ in this model. In addition, although the parameter 𝛿 remained adjusted for Convalescent group, the modeling of antibody dynamics allowed to better explain inter-individual variability of the elimination rate of infected cells compared to the viral model adjusted for group effects with an increase of 21% of explained variability. At the opposite, the adjustment of the viral model for mechanistic CoP (see Appendix B in S1 Appendix) led to a decrease of 21% of explained variability on 𝛿 compared to group effects. To quantify the benefit of modeling antibody dynamics, instead of using them as time-varying covariates in the viral model, we compared these two models in their quality to fit viral data using the root mean squared error (RMSE). As shown in Table 5, these two models integrating immune responses provided better ability of fitting viral data (i.e., lower RMSE) than the viral model adjusted for group effects. In addition, for all types of viral loads, the joint model highlighted better RMSE than the viral model with viral adjusted for immune markers. In accordance with RMSE results, the model well fitted viral load and antibody dynamics of all animals, with most of observations falling with the 95% individual prediction intervals. Individual fits of genomic viral load and antibody dynamics of eighteen NHPs randomly selected (three per group) are available in Figure S6 and Figure S7. In addition, we verified the ability of the model to capture the kinetics and the variability of the different observations across the entire NHP population by examining the Visual Predictive Check (Figure S8). Finally, the robustness of model parameter estimation was validated through a convergence assessment approach (Figure S9 and Figure S10).

**Table 5.** Root mean squared error calculated for the viral models, either with 𝜷 adjusted for group effects or mCoP, and the joint model

| Viral Loads | Viral Model with group effects<sup>1</sup> | Viral Model with mCoP<sup>2</sup> on $\beta$ and Conv effect on $\delta$ | Joint Model |
| --- | --- | --- | --- |
| gRNA in Nasopharynx | 0.702 | 0.686 | 0.660 |
| sgRNA in Nasopharynx | 0.580 | 0.507 | 0.494 |
| gRNA in Trachea | 0.603 | 0.630 | 0.613 |
| sgRNA in Trachea | 0.580 | 0.567 | 0.514 |

mCoP, mechanistic correlate of protection.
<sup>1</sup>Viral model with the viral infectivity $\beta$ adjusted for CD40.RBDv-Conv, CD40.PanCoV-Conv and mRNA-Conv groups, and the elimination rate of infected cells $\delta$ adjusted for Convalescent group.
<sup>2</sup>Viral infectivity $\beta$ is adjusted for the two immune markers as time-varying covariates, bAb(t) and nAb(t).

## Quantification of protective thresholds ACE2/RBD binding inhibition marker

### Evaluation of the protective threshold according to immunological background

Once the estimation of the model validated, estimated parameters were used to evaluate the within-host basic reproduction number ℛ_0_in the distinct groups. As shown in Figure 7B, animals with hybrid immunity presented a ℛ_0_ below 1 (mean [CI_95%_]: 2.0 × 10^―2^ [0.28; 0.83] × 10^―2^ for CD40.RBDv vaccine; 0.4 × 10^―2^ [0.023; 1.8] × 10^―2^ for CD40.PanCoV vaccine; 0.08 × 10^―2^ [0.0083; 0.3] × 10^―2^ for mRNA vaccine) meaning that NHPs were already protected against the propagation of the infection by their humoral responses at exposure. At the opposite, low levels of antibodies observed in Naive, only vaccinated and Convalescent animals at exposure, reflected by ℛ_0_ above 1 (mean [CI_95%_]: 5.0 [3.4; 7.9] for Naive; 3.0 [1.30; 4.20] for CD40.RBDv-Naive; 2.0 [1.30; 3.70] for Convalescent NHPs), made them susceptible to infection spreading. However, as shown in Figure 7C, representing the evolution of the effective reproduction number ℛ over time, the trigger and establishment of a strong neutralizing antibody response in vaccinated and convalescent animals allowed to control infection in the days following exposure. Based on these results, we then looked at the threshold of antibodies neutralizing ACE2/RBD binding requested at exposure to directly control within-host infection. We identified distinct protective thresholds for NHPs according to their immunological backgrounds. Considering the high variability of the MSD test for low values and the limit of positivity for nAb identified at 0.78 AU/mL (Table S1), a value lower than 1 AU/mL was predicted as protective for 95% of NHPs in mRNA-Conv and CD40.PanCoV-Conv groups by the model. Conversely, higher levels were requested in other groups with minimum levels of nAb ranging from 2.93 AU/mL in CD40.RBDv-Conv to 20.2 AU/mL in Naive groups at exposure. Figure 7A highlighted the consistency of these results with values of neutralizing antibody concentration observed over time in animals, with unprotected animals at the time of infection presenting ACE2/RBD binding inhibition levels lower than protective thresholds identified in their group.

**Figure 7.**
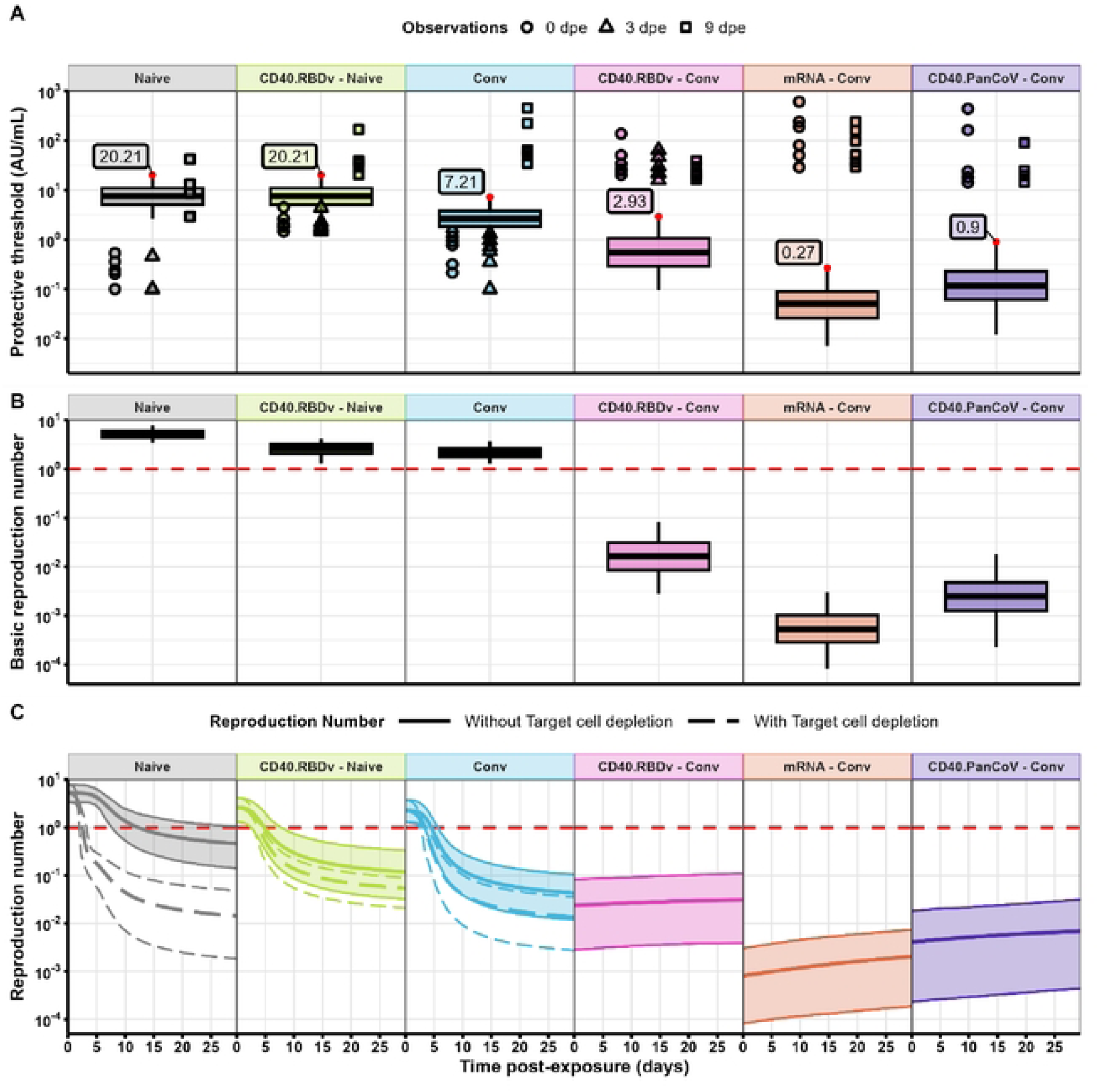
Evaluation of the protective threshold. 𝝉_𝒏𝑨𝒃_**and the reproduction number. (A)** Prediction of the ACE2/RBD binding inhibition protective threshold in each NHP group. Boxplots represent the distribution of the protective threshold predicted by the model (95% prediction interval). Red dots and labels highlight values of immune marker at which 95% of animals are protected against viral replication. Symbols represent ACE2/RBD binding inhibition data observed at 0 (circles), 3 (triangles) and 9 (squares) days post-exposure. (B) Prediction of the basic reproduction number ℛ_0_ in each NHP group. Boxplots represent the distribution of reproduction number predicted by the model (95% prediction interval). The horizontal red dashed line highlights the threshold ℛ_0_ = 1. (C) Evolution of the reproduction number over time in each NHP group. The dashed and solid thick lines represent the mean reproduction numbers predicted by the model with (i.e. effective reproduction number) and without (i.e., ℛ(𝑡), with 𝑇 = 𝑇_0_ ∀𝑡) uninfected target-cell depletion, respectively. White and shaded areas represent their respective 95% prediction intervals. The horizontal red dashed line highlights the threshold ℛ = 1.

### Efficient memory responses reduce the level of ACE2/RBD binding inhibition remaining protective against infection spreading

We took the opportunity provided by mechanistic models to explore counterfactual scenarios to better understand and interpret the distinct values of protective threshold found according to immunological background and their impact on viral control. To this end, we simulated viral and antibody dynamics under the counterfactual statement "What would have been viral control if similar levels of neutralizing antibodies had been observed in all groups at exposure ?" One interpretation of this scenario is the evaluation and comparison of viral control between groups if we had waited enough time after the last immunization in each of them to reach a given nAb level. When this initial antibody concentration was fixed at a low value equals to the protective threshold found in CD40.PanCoV-Conv group (i.e. 0.90 AU/mL, Scenario LownAb), as expected, viral control was only observed in mRNA-Conv and CD40.PanCoV-Conv animals (Figure S11 and Figure S12) highlighted by undetectable subgenomic RNA dynamics. At the opposite, nAb at exposure remaining lower than protective threshold in Naive, CD40.RBDv-Naive and Conv, a lack of viral control was still identified. As displayed in Figure 8 and Figure S12, showing how each hypothetical scenario impacted viral dynamics in each NHP group summarized by the five viral dynamics descriptors, the similar descriptors predicted by the model for the real and LownAb scenarios (e.g. peak VL or AUC) validated this hypothesis. The higher counterfactual viral replication (higher peak and AUC values) predicted in CD40.RBDv-Conv group compared to the real dynamics can be explained by the larger difference in antibody levels at exposure between factual and counterfactual scenarios than in other groups, and validated the necessity of a strong antibody response at exposure to efficiently control infection spreading (Figure 8 and Figure S12). While the same antibody levels and SARS-CoV-2 Delta infections were considered in all groups, these differences in viral control predicted between groups resulted from the distinct dynamics of ACE2/RBD binding inhibition observed after infection. As shown in Figure 9, faster increases of antibody responses were predicted in animals with hybrid immunity compared to others, in which a few days were required before the activation of an efficient immune response resulting in control of viral replication. Accordingly, these results pointed out the impact of memory immune response in the establishment and maintenance of a fast, efficient, and specific antibody response. The consideration of a higher initial concentration of neutralizing antibodies at exposure, with for instance nAb_0_ equals to 24 AU/mL (Scenario MidnAb), allowed to rapidly control viral replication in all groups, as highlighted by the significant reduction of VL peak or AUC values (Figure 8 and Figure S12) and resulting in undetectable sgRNA (Figure S11 and Figure S13), even in naive animals (Median [75%CI], -2.77 [-2.90; -2.55] log_10_ copies/mL for gRNA peak and -4.34 [-4.72; -3.97] log10 copies/mL for sgRNA peak; -1.22 [-1.44; -1.03] × 10^8^ copies.day/mL for gRNA AUC and -8.97 [-12.1; -6.59] × 10^6^copies.day/mL for sgRNA AUC) despite their lack of efficient B cell immune response. This control is explained by the level of antibodies at exposure higher than the protective threshold, resulting in a rapid blockage of new infection (reduction of sgRNA viral peak, and acute and clearance stages duration). Nevertheless, the lack of efficient memory immune response in naive animals is translated into a potential increase of genomic viral load acute and clearance stages duration (Figure 8) induced by a significant reduction of uninfected target-cell depletion, which drives most of viral control in naive animals (see Figure 7C). Accordingly, these results validate that a strong and efficient immune response at exposure is not only driven by a high concentration of neutralizing antibodies but also by their efficacy to neutralize viruses. Finally, for all groups, complete viral control is observed in the case of a high initial level of nAb (Scenario HinAb; Figure S14).

**Figure 8.**
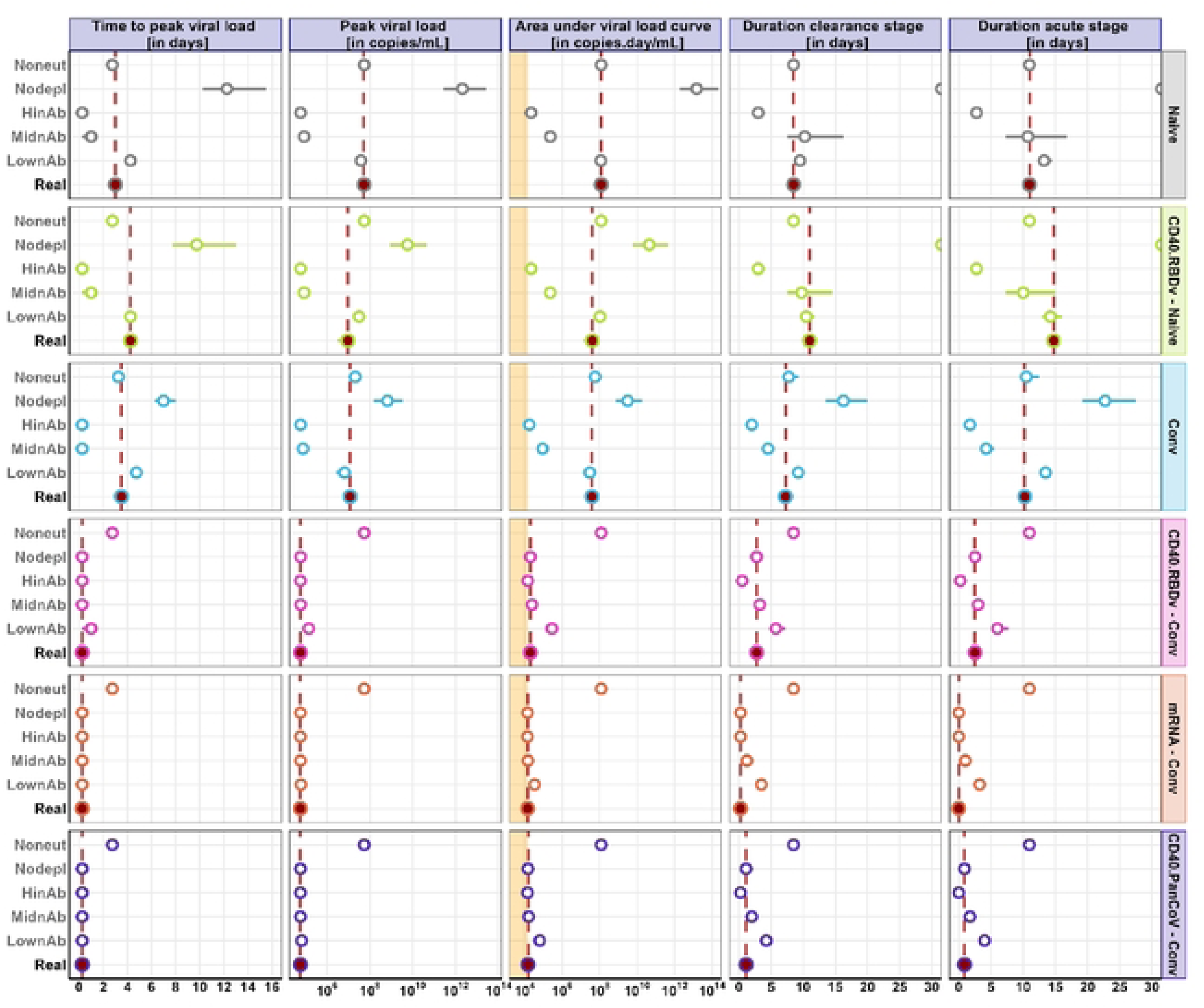
Viral dynamics descriptors of genomic RNA (gRNA) predicted in real and all counterfactual scenarios. Comparison of the five viral descriptors (Time to peak, Peak, AUC, Clearance and Acute stages duration) predicted in each NHP groups in 5 five counterfactual/hypothetical scenarios and the real/factual scenario (y-axis) to validate modeling hypothesis. *<u>LownAb</u>:* nAb at exposure fixed at a low value in all groups, 𝑛𝐴𝑏_0_ = 0.90 AU/mL; *<u>MidnAb</u>*: nAb at exposure fixed at a medium value in all groups, 𝑛𝐴𝑏_0_ = 24 AU/mL; *<u>HinAb</u>*: nAb at exposure fixed at a high value in all groups, 𝑛𝐴𝑏_0_ = 437 AU/mL; *<u>Nodepl</u>*: no depletion of uninfected target-cells after Delta infection in all groups, 𝑇(𝑡) = 𝑇_0_; *<u>Noneut</u>:* nAb are non-efficient to neutralize viruses in all groups, 𝜀_𝐴𝑏_(𝑡) = 0. Dots- and horizontal-colored segments represent the mean and the 95% prediction intervals of predicted descriptors. Vertical red dashed lines represent the mean value predicted in the real scenario. Colored shaded area highlights range of descriptors (Peak or AUC) values estimated only with undetectable viral measures.

**Figure 9.**
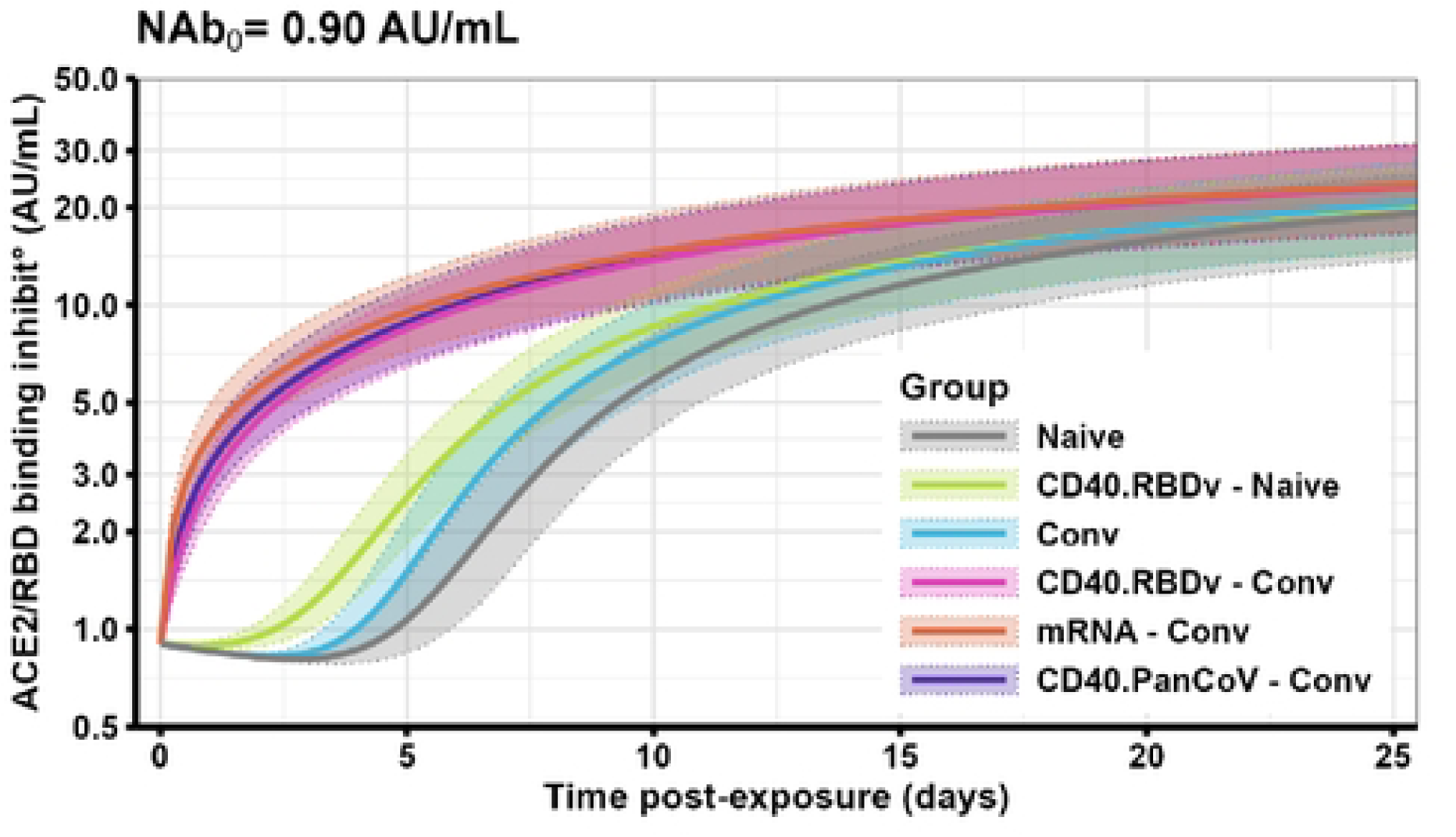
What if the neutralizing antibody concentration at exposure in all groups had been equal to a low value of 0.90 AU/mL [Counterfactual Scenario LownAb]. Predicted dynamics of antibodies neutralizing ACE2/RBD binding that would have been observed after infection in each group of NHP. Thick lines and shaded areas represent the mean dynamics and the 95% prediction intervals, respectively, for Naive (in gray), CD40.RBDv-Naive (in light green), Convalescent (in blue), CD40.RBDv-Conv (in pink), CD40.PanCoV-Conv (in purple) and mRNA-Conv (in orange) NHP groups.

In conclusion, these counterfactual scenarios allowed to explain the differences in antibody levels neutralizing ACE2/RBD binding considered as protective in the distinct NHP groups as an effect of the efficacy of memory immune response to elicit fast and specific antibody response.

### Viral control driven by target-cell depletion in naive immune systems and by humoral responses in immunized animals

The humoral immune response is not the only mechanism impacting the control of viral dynamics after infection. Indeed, the depletion of uninfected target cells also appeared as a key element in naive animals. To distinguish the implications of these two mechanisms, we studied two counterfactual scenarios: one assuming the absence of target-cell depletion (scenario Nodepl) and the second considering inefficient antibody response (scenario Noneut). In both cases, the ability to contain viral infection was compared to a real scenario.

As shown in Figures 8, 10A, and S11, viral dynamics were not impacted by target-cell depletion in the case of hybrid immunity. A moderate effect was found in CD40.RBDv-Naive and Convalescent groups characterized by a small increase of the time of gRNA peak (+1.23 [0.25; 2.75] days in convalescent and +1.01 [0.00; 4.00] days in vaccinated NHPs), of the duration of clearance stage with +2.04 [0.00; 5.50] days in convalescent and +2.62 [0.00; 9.75] days in vaccinated NHPs, and of acute stage with +3.26 [0.50; 8.25] days in convalescent and +3.43 [0.00; 12.30] days in vaccinated NHPs. Conversely, a high loss of viral control was observed in naive animals with a significant increase of the time of gRNA peak (+10.68 [5.00; 26.75] days) and peak values exceeding 10^10^ copies/mL, highlighting the inefficacy of immune system to control viral replication within the 30 days following infection.

The removal of an efficient antibody response in the different groups highlighted the strong impact of these latter on viral dynamics in animals with hybrid immunity, as shown by the uncontrolled dynamics observed in Figure 10B. On the contrary, no modification of dynamics was induced in Naive animals, while a moderate effect of immune responses on viral dynamics was identified in both CD40.RBDv-Naive and Convalescent groups. In conclusion, we identified the humoral immune response as the major immune mechanism controlling viral replication in immunized NHPs, while control of viral infection was mostly driven by the reduction of available virus target-cells over time in the case of naive immune systems. In an in-between regime, both mechanisms were responsible for control in only vaccinated or convalescent animals.

**Figure 10.**
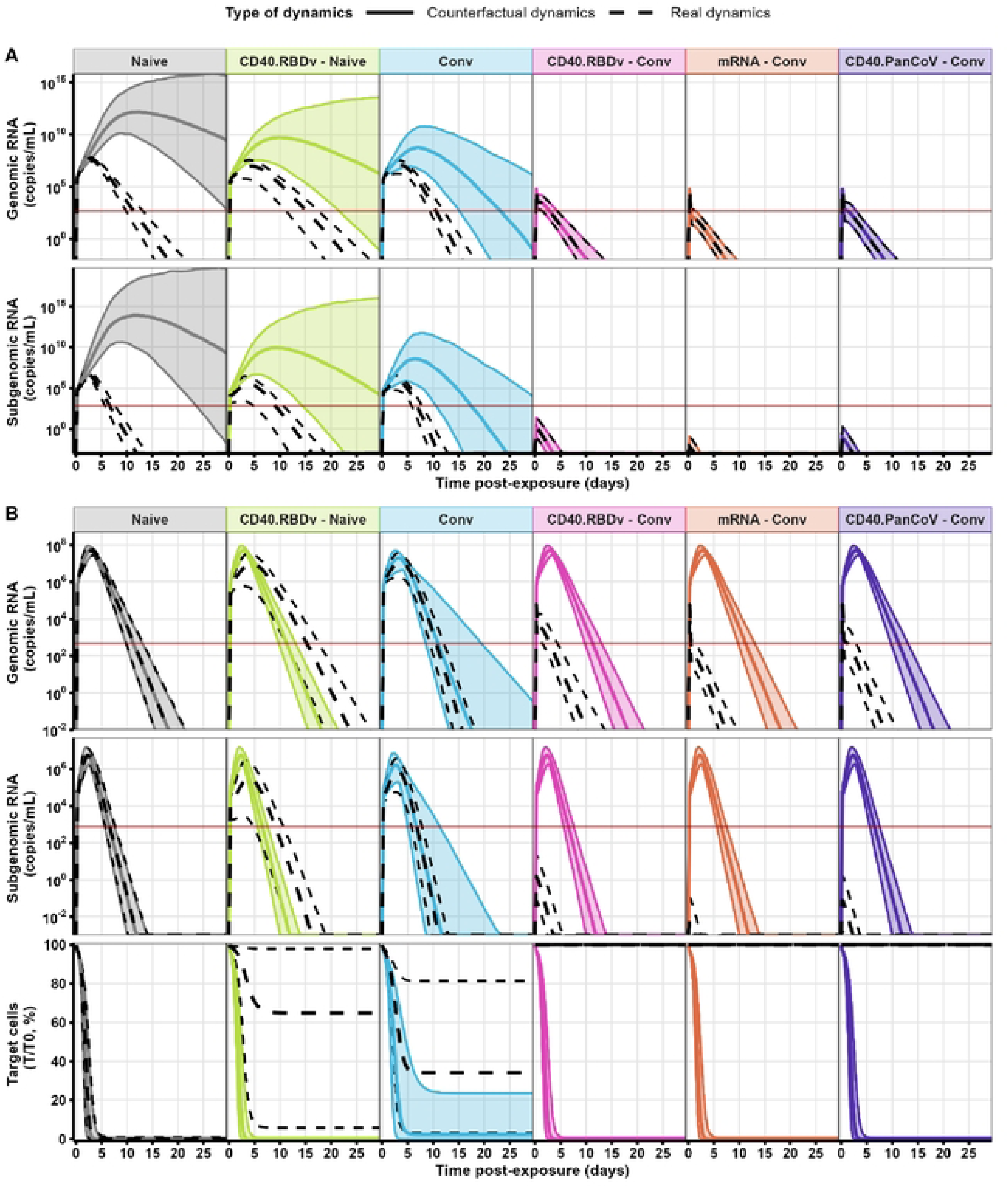
Impact of the absence of uninfected target-cell depletion or efficient immune responses on the control of viral replication after infection. (A) Absence of uninfected target-cell depletion [Counterfactual scenario Nodepl]. (B) Absence of efficient immune responses [Counterfactual scenario Noneut]. (A-B) Comparison of real viral dynamics predicted by the model and dynamics predicted by counterfactual statements in each NHP group. Colored thick lines and shaded areas represent mean dynamics and 95% prediction intervals predicted obtained in counterfactual scenario. Dashed lines represent mean dynamics and 95% prediction interval obtained in real scenario. Horizontal solid red lines highlight viral load limit of detection, with LOD = 476 copies/mL for genomic RNA and 749 copies/mL for subgenomic RNA.

## Discussion

In this original modeling work, we proposed a mechanistic model to jointly describe post-infection viral and antibody dynamics, and their mutual interactions, with the main purpose to quantify protective thresholds of neutralizing antibodies allowing rapid viral control after SARS-CoV-2 infection. We estimated the model on preclinical data, including viral loads, binding, and neutralizing antibody kinetics against SARS-CoV-2 Delta variant collected in a large number of NHPs with various immunological backgrounds: non-vaccinated or vaccinated naive, and non-vaccinated or vaccinated (with CD40.RBDv, CD40.PanCoV, or mRNA vaccines) convalescent animals from previous Wuhan infection. We first confirmed results from our previous modeling work [32] regarding 1) the main immune mechanisms induced by vaccination and/or natural immunity impacting viral control following viral challenge, and 2) the mCoP capturing these effects. In particular, we showed that antibodies generated after immunization and inhibiting ACE2/RBD binding are reliable mechanistic CoPs capturing the effect of the immune system to block new infection of susceptible target cells. Moreover, we found an additional effect of natural immunity besides neutralizing antibodies, promoting the elimination of infected cells, but without identifying potential mCoPs. The extension of the model for antibody dynamics and their neutralizing functionality led to the identification of vaccine- and natural infection-induced mechanisms of action of humoral responses. We found that B cells and plasma cells responses, all captured by a single ODE compartment here, were strongly dependent on NHP immunity profiles, highlighted by 1) a higher concentration of immune cells in vaccinated or hybrid animals at exposure, and 2) a faster establishment of immune responses in convalescent animals, regardless vaccinations status. The model also provided evidence of a stronger antibody production by plasma cells after a single immunization event, whether from primary vaccination or previous Wuhan infection, although the latter appeared more efficient at neutralizing viruses in cases of hybrid immunity. These results are in accordance with neutralizing antibodies with higher affinity and specificity as expected in the case of multiple timely spaced antigen contacts [79] highlighting the strong benefit of hybrid immunity to induce a faster and more protective immune response after infection. Through this modeling work, we could derive minimum protective neutralizing antibody concentrations leading to rapid viral control. Distinct thresholds identified for the different groups of animals pointed out the key role of B-cell memory responses in the prediction of the post-vaccination and infection responses [79].

The analysis conducted in this paper strongly improved our understanding of the causal impact of humoral immune responses on the control of viral dynamics. Nevertheless, these results accentuated the necessity to evaluate CoPs as dynamical immune markers instead of focusing on specific time points as usually done in the literature [23,24,26]. Indeed, while nAb levels were similar between naive and convalescent NHPs at exposure, better viral control was observed in immunized animals, resulting from the mobilization of their antibody and B-cell memory responses. Moreover, the distinct thresholds evaluated in CD40.RBDv-Naive and Convalescent NHPs sustained the fact that CoPs may differ for immunities induced by infection or vaccination [8]. In that context, one strong limitation of our model is the integration of the immunological background of NHPs as external information to explain post-Delta infection antibody dynamics. Modeling the overall process from the first Wuhan infection could contribute to overcome issues encountered due to the simplicity of the model. Considering of a more complex model for humoral responses should better explain the immune mechanisms behind the observed saturation level of antibodies in hybrid immune systems and thus improve understanding of the identified immunological background effects. The nonsignificant effect of hybrid immunity on the antibody production rate by ASCs may result from a severely constrained model. For instance, Dari et al. [49] proposed a model describing B cell responses after recognition of viral antigens, including germinal centers and memory B cells, their respective generation of long-lived and short-lived plasma cells, and the resulting production of antibodies. However, estimating this model requires data with a longer post-infection follow-up to study the longevity of the humoral immune response.

The next question beyond quantifying a valid mCoP is its interpretation in the actual context of viral evolution [8]. In fact, the protective threshold predicted here involved only the Delta variant, whether as the strain of infection or the strain targeted by antibodies. However, with the emergence of Omicron subvariants and their ability to escape immunity, these thresholds may no longer be appropriate. Similarly, the same concentration of neutralizing antibodies targeting the ancestral strain may lead to lower viral control of Delta infection [31]. Therefore, including notions such as cross-reactivity, or at least cross-variant neutralization, in the modeling may help derive variant-specific protective thresholds against a particular strain of infection. Furthermore, as strongly discussed in the recent literature, a better understanding of how immune imprinting can impact SARS-CoV-2 vaccine-induced responses against new variants is crucial for developing and evaluating next-generation vaccines, such as CD40-targeting vaccines or variant-specific mRNA vaccines. In the case of pre-Omicron variants, and as demonstrated by this work, immune imprinting has not necessarily been shown to compromise protection, reinforcing the benefit of hybrid immunity in inducing a strong immune response [80,81]. However, the effects of imprinting remain unclear for more recent variants. On one hand, the FDA and some studies [81,82] recommended formulating new mRNA vaccines with a single monovalent Omicron spike protein to improve variant-specific immunogenicity and reduce the influence of prior immunization with ancestral vaccines. On the other hand, other studies [82,83] have demonstrated the benefit of recalling B cell responses against the ancestral strain to enhance neutralization activities. While neutralization titers have been identified as reliable CoP for all pre-Omicron variants, extensive mutations in the RBD acquired in Omicron variants have enhanced their capacity to escape neutralizing antibodies [2], thus significantly reducing the protective effect of neutralizing antibodies. However, as pointed out by Clark et al. [21] or Zhang et al. [84], anti-spike non-neutralizing antibodies, which can bind to a larger part of the Spike protein and have already been identified as reliable CoP [23,24], are less impacted by Omicron mutations and can still induce Fc-dependent effector functions, contribute to viral control, and mediate protection against Omicron, in combination with anti-spike T-cell responses. Extending our model to integrate Fc-dependent effector functions on viral dynamics is relevant in this global immunological context, focusing in particular on the enhancement of virus and infected cells clearance [60,65], as well as T-cell immune responses [58]. Of note, the two CD40-based vaccines tested in this study were designed to incorporate key mutations conserved across SARS-CoV-2 variants, in addition to a conserved nucleocapsid sequence rich in T-cell epitopes. The cross-reactivity against VoC will be evaluated in ongoing clinical trials testing these two candidate vaccines.

Similarly, the research of mCoP, able to capture the effect of natural immunity on the enhanced elimination of infected cells, could strongly benefit from the consideration of the functions of non-neutralizing antibodies. In fact, this latter may result from the antiviral functionality of non-neutralizing antibodies, which can mediate protection by activating immune effector cells via the binding to their Fc receptors, thereby inducing Fc-dependent effector functions such as antibody-dependent cellular cytotoxicity (ADCC), antibody-dependent cellular phagocytosis (ADCP), or modulation of T cell responses [21,84,85].

The role of ADCC in natural infection has been previously proven [85,86]. Therefore, measuring these non-neutralizing antibody functions could capture this additional effect on infected cell clearance. However, this requires collecting a large number of variables, which is more complex than standard binding and neutralization assays [84]. Furthermore, since this effect was observed only in convalescent NHPs in the absence of vaccination, a major limitation of our study that may need to be addressed in future analyses is the small sample size within each group of animals (n ≤ 6 per group), which restricts the power to identify group-specific effects.

## Acknowledgments

We thank Emma Burban, Benoit Delache, Sebastien Langlois, Joanna Demilly, Nina Dhooge, Pauline Le Calvez, Victor Magneron, Eléana Navarre, Maxime Potier, Jean-Marie Robert, Quentin Sconosciuti and the team of Animalliance / Uniivo for the NHP experiments; Loïc Pintoree, Maxence Galpin-Lebreau and Julie Morin for the RT-qPCR and for the preparation of reagents; Léo Joffroy, Julien Dinh, Alexandre Baillet and Elodie Guyon for the NHP sample processing. The virus stocks were obtained from the Biodefense, Research Resources, and Translational Research branches of the NIH/NIAID/DMID/OBRRTR/RRS (Program Officer, Clint Florence, Ph.D.).

This work was supported by INSERM and the Investissements d’Avenir program, Vaccine Research Institute (VRI), managed by the ANR under reference ANR-10-LABX-77-01 and the PSPC COVID-19 – Projet EVIDENCE funded by the BPI (Banque Publique d’Investissement). The Infectious Disease Models and Innovative Therapies (IDMIT) research infrastructure is supported by the “Programme Investissements d’Avenir”, managed by the ANR under reference ANR-11-INBS-0008. The Fondation Bettencourt Schueller and the Region Ile-de-France contributed to the implementation of IDMIT’s facilities.

Figure 1 and Figure 2 were created with BioRender.com. We thank Simulations Plus, Lixoft division for the free academic use of the MonolixSuite. Numerical computations were in part carried out using the PlaFRIM experimental testbed, supported by Inria,

CNRS (LABRI and IMB), Université de Bordeaux, Bordeaux INP, and Conseil Régional d’Aquitaine (see https://www.plafrim.fr).

## Author Contributions

**Conceptualization:** MA, RM, RLG, RT, YL, MP

**Data curation:** LB, MC, NDB **Formal analysis:** MA **Funding acquisition:** MC, YL

**Investigations** MA, RM, RLG, RT, YL, MP, MC **Methodology:** MA, RM, RLG, RT, YL, MP **Resources:** FR

**Supervision:** MP, RT, YL

**Software:** MA

**Validation:** MA, RM, RLG, RT, YL, MP

**Visualization:** MA, RM

**Writing - original draft:** MA, MP

**Writing - review & editing:** All

## Conflict of interest statement

The remaining authors declare no competing interests.

## Data and code availability

Data from this study are available on the secured data repository platform Labkey of whom access can be asked upon reasonable requests to the corresponding authors.

The original code (mlxtran models and R codes) developed in this work is available and free-of-cost on github (Inria SISTM Team) in the folder ‘EarlyViralandAntibodyDynPostInfection’ at the link https://github.com/sistm/SARSCoV2modelingNHP.

## Supporting information

**Supplementary Figures – Fig. S1 to S14 Supplementary Tables – Table S1 Supplementary Files – Appendix A to D**

## Notes

### Competing Interest Statement

The authors have declared no competing interest.

